# Release of the pre-assembled naRNA-LL37 composite DAMP re-defines neutrophil extracellular traps (NETs) as intentional DAMP webs

**DOI:** 10.1101/2022.07.26.499571

**Authors:** Francesca Bork, Carsten L. Greve, Christine Youn, Sirui Chen, Yu Wang, Masoud Nasri, Jule Focken, Jasmin Scheurer, Pujan Engels, Marissa Dubbelaar, Katharina Hipp, Birgit Schittek, Stefanie Bugl, Markus W. Löffler, Julia Skokowa, Nathan K. Archer, Alexander N.R. Weber

## Abstract

Neutrophil extracellular traps (NETs) are a key antimicrobial feature of cellular innate immunity mediated by polymorphonuclear neutrophils (PMNs), the primary human leukocyte population. NETs trap and kill microbes but have also been linked to inflammation, e.g. atherosclerosis, arthritis or psoriasis by unknown mechanisms. We here characterize naRNA (NET-associated RNA), as a new canonical, abundant, and unexplored inflammatory NET component. naRNA, upon release by NET formation, drove further NET formation in naïve PMN, and induced macrophage and keratinocyte activation via TLR8 in humans and Tlr13 in mice, in vitro and in vivo. Importantly, in vivo naRNA strongly drove skin inflammation, whereas genetic ablation of RNA sensing drastically ameliorated psoriatic skin inflammation. Rather than accidentally assembling with LL37 on the NET, naRNA was intracellularly pre-associated in resting neutrophils as a ‘composite DAMP’, thus highlighting NET formation as a DAMP release process. This re-defines sterile NETs as an intentionally inflammatory agent, signaling and amplifying neutrophil activation. Moreover, in the many conditions previously linked to NETs and extracellular RNA, TLR-mediated naRNA sensing emerges as both potential cause and new intervention target.

**Graphical abstract:** 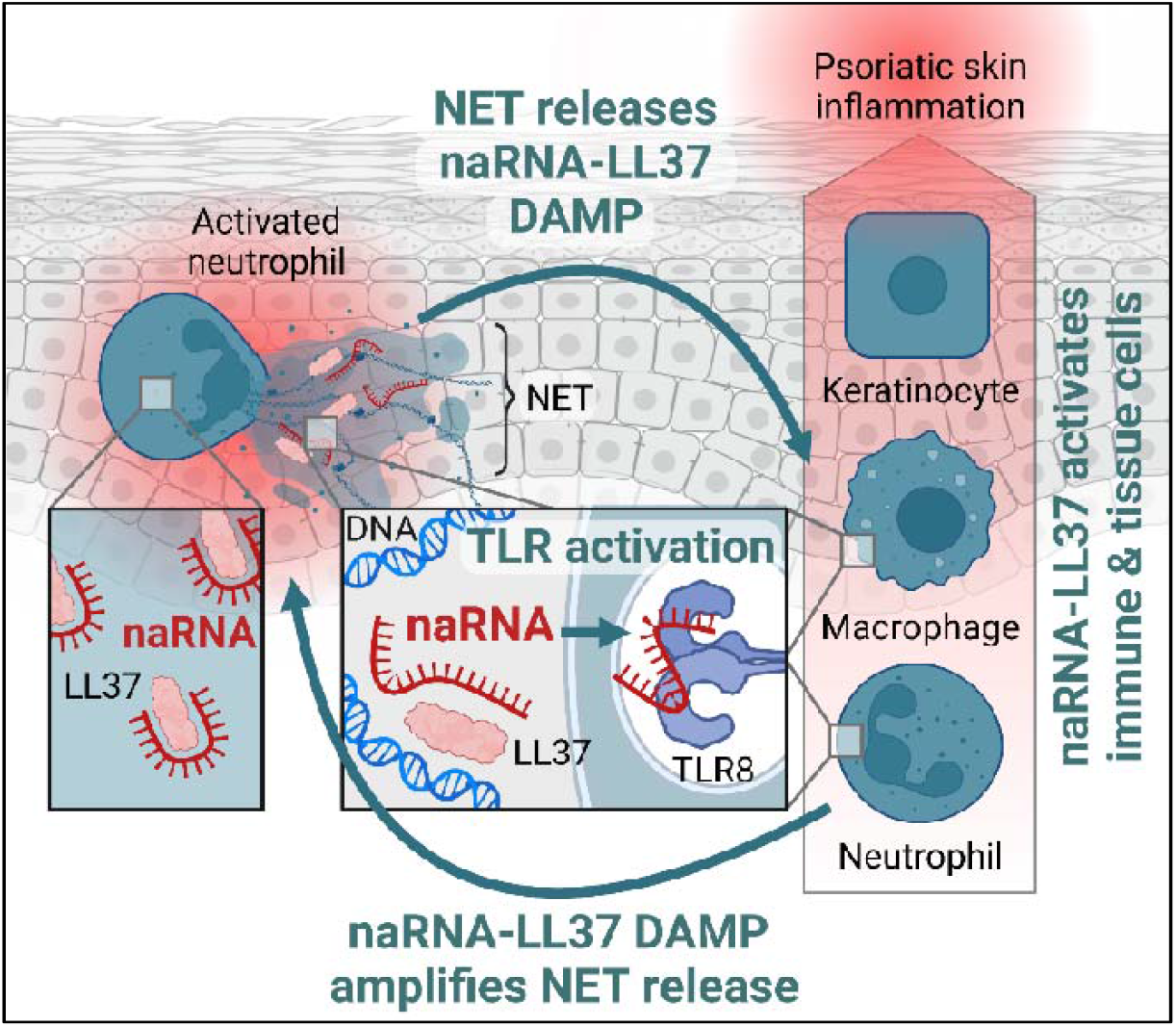

Created with biorender.com

## Introduction

The formation of Neutrophil Extracellular Traps (NETs), since its discovery in 2004 (*1*), has emerged as a fascinating phenomenon of host defense. Hereby neutrophils, the primary leukocyte population, extrude their genomic DNA to form web-like structures that, similar to barbed wire roadblocks, trap and kill microbial invaders such as bacteria or fungi (*2*). DNA is thus a defining structural and functional feature of NETs. Additionally, DNA-associated proteins, histones and HMGB-1, antimicrobial peptides like LL37 and enzymes such as myeloperoxidase (MPO) are released during the formation of NETs and contribute to their antimicrobial function (*2*). With their primary role firmly established to be antimicrobial, the reported role of NETs in sterile inflammatory conditions such as atherosclerosis (*3*), rheumatoid arthritis (RA) (*4*) and COVID-19 (*5*) has remained somewhat enigmatic, but generally an accidental phenomenon. Specifically, the observed inflammation-promoting effects of sterile NETs have not been mechanistically linked to a specific component of NETs, despite the fact that >1,000 publications on NETs have sought to detail the multi-faceted phenomenon and the processes that lead to its execution (*6*). However, with the main focus devoted to DNA and protein components, another primary cellular biomolecule, RNA, has so far received very little attention in vertebrates. Interestingly, a study in insects showed that hemocytes (macrophage-like immune cells) can release both DNA and RNA in NET-like structures during microbe-triggered clotting reactions, and in response to extracellular RNA or DNA (*7*). Our lab recently provided evidence of a responsiveness of human polymorphonuclear neutrophils (PMNs) to RNA, but not DNA, in combination with the antimicrobial peptide LL37 (*8*). In addition, earlier work described a role for DNA- and RNA-LL37 complexes in the activation of plasmacytoid dendritic cells (*9, 10*). This has given rise to the notion that under certain conditions tolerance to self-DNA and RNA ‘accidentally’, e.g. in psoriasis patients, can be broken by LL37 (*11*), involving the nucleic acid-sensing TLR7, TLR8 (ssRNA) and TLR9 (DNA) in humans, and Tlr7, Tlr13 and Tlr9 in mice (*12, 13*). In general, many roles have been ascribed to so-called extracellular RNA (exRNA), for example, macrophage polarization, recruitment of leukocytes to the site of inflammation, leukocyte rolling at the vascular endothelium, as well as integrin-mediated firm adhesion of immune cells and promotion of thrombosis (*14*). Additionally, exRNA is considered as a reliable biomarker for various diseases such as cancer or cardiovascular pathologies (*15, 16*). However, RNA being considered a short-lived biomolecule, physiological sources of exRNA have been somewhat unclear. Under sterile conditions, vascular injury, tissue damage, or ischemia have been suggested to trigger release of exRNA along with other cellular material (*14*) but it remains unclear whether these relatively slow processes could amount to physiologically detectable exRNA amounts in the serum of patients, or whether a rapid, as-yet unidentified process of RNA extrusion would have to be postulated.

We show here that RNA contained in NETs, so-called NET-associated RNA or naRNA, drives such a self-amplifying inflammatory loop not only engaging PMNs but also macrophages and keratinocytes. naRNA responsiveness was dependent on *TLR8* and *Tlr13* in human and murine myeloid immune cells, respectively. Notably, in mice naRNA caused considerable skin inflammation in a Tlr13-dependent manner and in a well-established model of psoriatic skin inflammation, genetic ablation of RNA sensing strongly ameliorated skin inflammation. Importantly, the observed pre-association of naRNA with LL37 in resting neutrophils from healthy donors indicates that NETs are strictly intended for extrusion of this novel ‘composite’ DAMP. Not only does this show that the presumed ‘breaking of tolerance’ by LL37 is non-accidental. As the majority of pathological conditions in which NETs have been described in are sterile, our data re-define the primary purpose of NET formation to be DAMP release, with naRNA-LL37 acting as a molecular beacon to report on PMN activation and alert tissue and other immune cells.

## Results

### naRNA is a common component of NETs

To first investigate whether naRNA was a common component of NETs, we compared NETs from human PMNs formed in response to the well-defined molecular agonists PMA, different complexes of LL37 with purified RNA (synthetic, as well as total fungal and bacterial RNA from *S. aureus* and *C. albicans*, respectively), nigericin and the live pathogen *C. albicans*. Confocal microscopy analysis using the well-characterized rRNA-specific antibody, Y10b (*17*), revealed the presence of RNA in all corresponding NETs, independent of stimulus and whether NET formation was ‘suicidal’ or not (*6*) (Fig. 1A, quantified in B; control in Fig. S1A): the RNA signal was readily detectable along the web-like DNA threads in human NETs. Similar results were obtained in NETs released by murine bone marrow-derived PMNs (BM-PMNs, Fig. 1C; control Fig. S1B). To avoid the use of staining reagents, we also metabolically labeled primary human hematopoietic stem cells (HSCs) with 5-ethynyluridine (5-EU), a nucleotide that can be incorporated in cellular RNA but not DNA, and that is amenable to labeling by click chemistry (*18*). These HSCs were then differentiated to neutrophils (*19*) and the differentiation validated by microscopy and flow cytometric analysis (Fig. S1C, D). NETs released from HSC-derived PMNs could indeed be labeled with click reagents conferring a fluorescent label only when grown in 5-EU-containing medium (Fig. 1D). Moreover, the overlap between click-label and rRNA confirmed the specificity of the staining with anti-rRNA antibodies and indicated that naRNA contained rRNA but also other types of cellular RNAs. This experiment also unequivocally confirmed that upon PMN stimulation, cellular RNA is turned into a component of the NET, wherein it decorates the fine, web-like DNA structures. The localization of naRNA on DNA strands was further confirmed by high-resolution fluorescence microscopy and 3D analysis (AiryScan, Fig. 1E and Supplemental Movie S1) as well as scanning electron microscopy (SEM, Fig. 1F; controls Fig. S1E). Extraction of naRNA from PMA-induced NETs (Fig. 1G) and subsequent RNA-sequencing analysis confirmed that purified naRNA contained multiple RNA types, with noticeably more non-ribosomal RNA than the corresponding total cellular RNA of PMNs (Fig. 1H). Collectively, these results indicate that naRNA is a canonical component of NETs released upon multiple stimuli from both human and murine neutrophils.

**Fig. 1:**
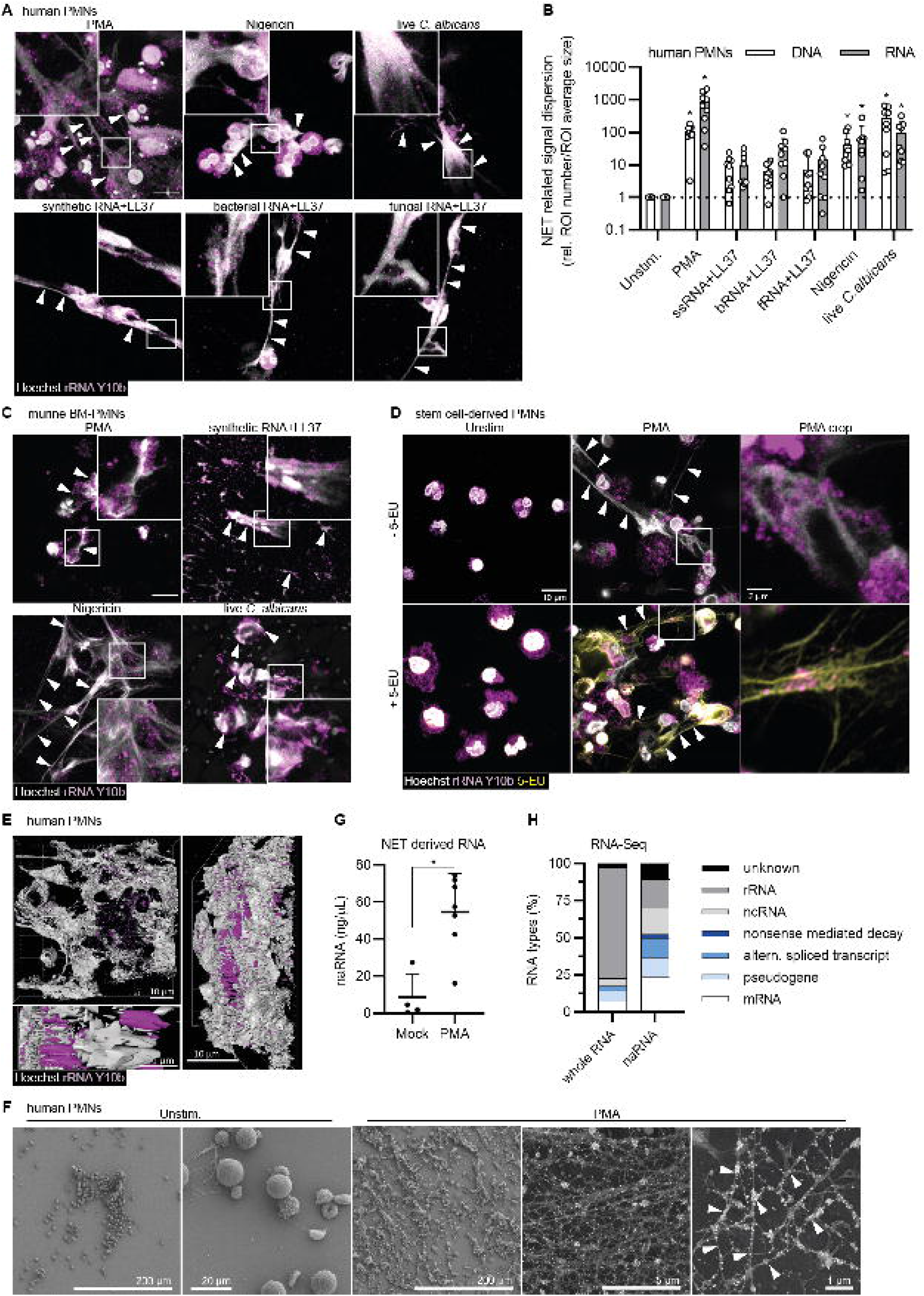
naRNA is a canonical component of NETs. **(A, B)** Confocal microscopy of primary human PMNs stimulated as indicated for 3 h and stained for naRNA (anti-rRNA Y10b, magenta) and DNA (Hoechst 33342, white, n = 3, scale bar: 10 μm, arrowheads point to selected NET strands; representative images in **A**) were quantified in **B** (each dot represents one image, *p<0.05 according to one-way ANOVA). **(C)** Confocal microscopy of primary murine BM-PMNs stimulated as indicated for 16 h and stained as in **A** (n = 3, representative images, scale bar: 10 μm). **(D)** Confocal microscopy of primary human stem cells differentiated *in vitro* with/without 100 μM 5-ethynyluridine (5-EU), click-labeled with a fluorescent dye (yellow, total RNA), anti-rRNA and Hoechst 33342 as in **A** (n = 3, representative images, scale bar: 10 μm, 2 μm in cropped image). **(E)** as in **A** but showing 3D image reconstruction of z-stacks (n = 3, representative images, scale bar: 10 μm). **(F)** Scanning electron microscopy of human primary PMNs treated as indicated and using anti-rRNA primary and immunogold (white arrow)-labeled secondary antibodies and silver enhancement (n = 3, representative images, scale bars as indicated; the two rightmost images show composite images with signals from secondary electron and backscattered electron detectors for topography and additional material information, respectively). **(G)** Agilent TapeStation quantification of naRNA isolated from mock or PMA NETs (from n = 4-6 donors, combined data, mean+SD, each dot represents one biological replicate/donor, *p<0.05 according to Mann-Whitney test). **(H)** RNAseq of PMA NET naRNA (n = 4) and whole PMN RNA (n = 1) (combined data).

### naRNA is a potent ‘composite’ DAMP propagating NET formation in primary PMNs

A primary function of NETs is host defense by trapping and killing bacteria. Therefore, we first explored if naRNA participated in this process (*1*). However, the antibacterial effect of NETs on live *Staphylococcus aureus* (evidenced by lower CFUs compared to resting PMNs) was similar to that of RNase-treated NETs, whereas DNase digestion reduced the drop in CFU discernibly but non-significantly (Fig. S2A). Consequently, we turned our attention to a possible role of naRNA as a DAMP, since primary PMNs can respond to synthetic RNA-LL37 complexes with NET formation, and our earlier analysis indicated that NETs also contain LL37 (*8*). To prepare a naRNA-containing stimulant, PMA-induced NETs were prepared as described in Methods and the original stimulus, PMA, removed by extensive washing before harvesting (Fig. 2A). Interestingly, these ‘PMA NETs’, when applied to naïve PMNs from another donor, potently induced new NETs (Fig. 2B, quantified in C, controls in Fig. S2B). Treatment with an RNase inhibitor enhanced the stimulatory effect of PMA NETs rendering a 1:500 dilution of PMA-NETs more effective than a 1:50 dilution of non-treated PMA-NETs (Fig. 2B, C, controls Fig. S2B). Interestingly, similar NET preparations from mock-treated PMNs (here referred to as ‘mock NETs’) did not stimulate NET formation under the same experimental conditions. To rule out the possibility that residual PMA in PMA NETs could be responsible for the observed stimulatory effects, RNase treatment was also performed which strongly reduced NET formation (Fig. 2D, quantified in E; controls Fig. S2C). The destruction of naRNA in NETs was further confirmed by rapid loss of the RNA signal (using the RNA-selective dye, SYTO RNAselect (*8*) in time-lapse digestion analysis (Supplemental Movie S2). Bearing in mind that PMNs do not respond to DNA or DNA-LL37 complexes (*8*), the opposing effects of RNase inhibitor and RNase treatments thus clearly indicated naRNA (and not DNA) to be the relevant immunostimulatory component for NETs to drive the activation of naïve PMNs. In line with our previous work showing that synthetic RNA or LL37 alone cannot trigger PMN activation (*8*), purified ‘naked’ (i.e. (stripped of DNA or proteins) naRNA isolated from PMA NETs (*cf*. Fig. 1G) was unable to activate NET formation; however, re-complexing with exogenous LL37 was sufficient to restore NET formation (Fig. 2F, controls in Fig. S2D). naRNA (*cf*. Fig. 2F) and LL37 (*8*) on their own are thus not DAMPs but, in the context of NETs, act in concert as a novel type of ‘composite’ DAMP.

**Fig. 2.**
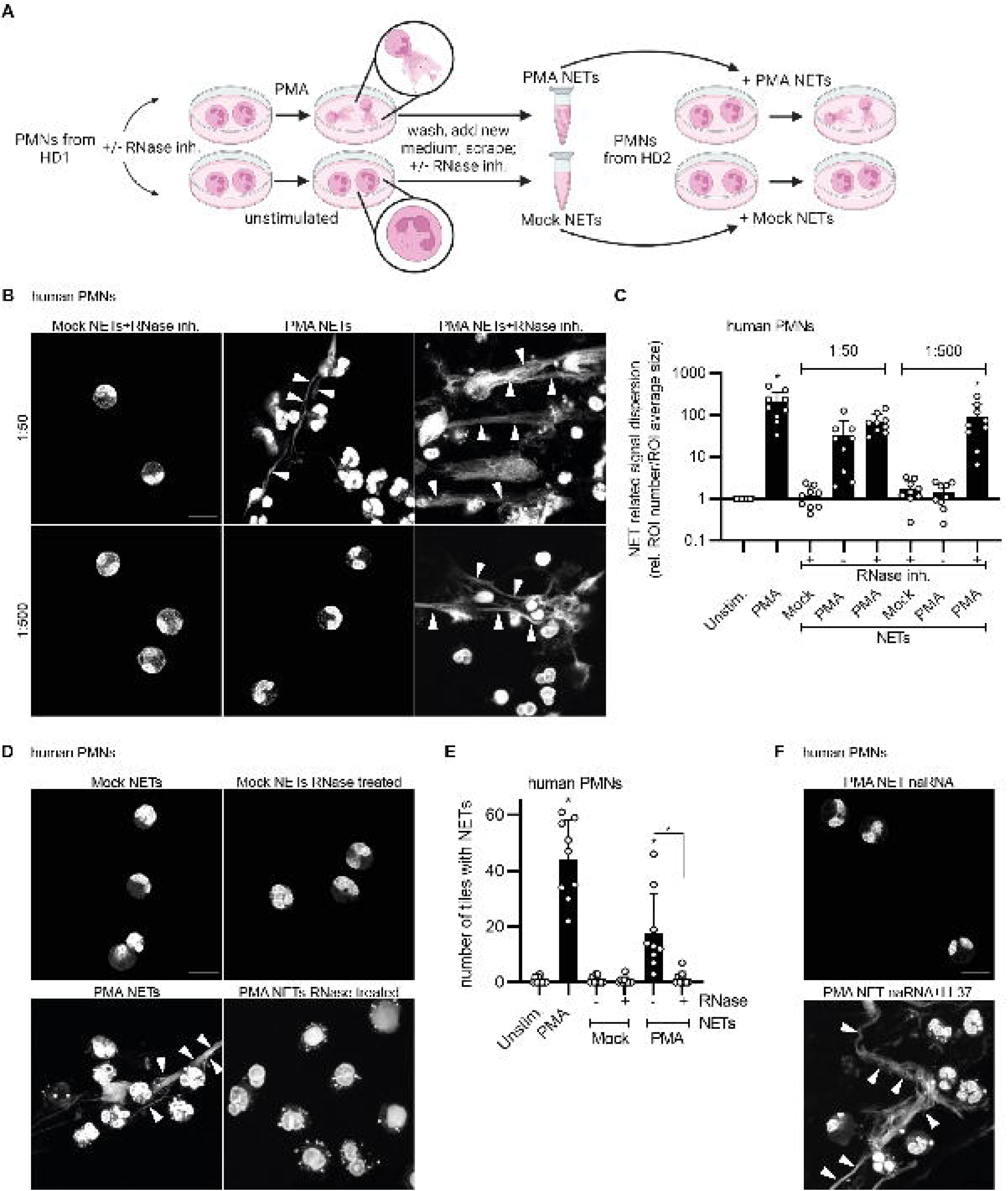
naRNA is a composite DAMP driving NET propagation in human PMN. **(A)** Workflow for NET content preparation from one donor and transfer to naïve human primary PMNs from a second donor. **(B)** Confocal microscopy of primary human PMNs stimulated with NET content harvested with/without RNase inhibitor and diluted 1:50 or 1:500, and then stained for NETs/DNA (Hoechst 33342, n = 9, representative images, scale bar: 10 μm). **(C)** Quantification of **B** using DNA (Hoechst 33342) signal to quantify NET formation (n = 3, combined data, mean+SD, each dot represents one image, *p<0.05 according to one-way ANOVA). **(D)** as in **B** but with/without pre-digestion of NET content with RNase A (n = 3, representative images, scale bar: 10 μm). **(E)** Quantification of **D** (n = 3, combined data, mean+SD, each dot represents the number of NET-positive tiles in one image quantified from three images/condition, *p<0.05 according to one-way ANOVA). **(F)** As in **B** but using purified naRNA (*cf*. Fig. 1G) alone or in complex with exogenously added LL37 (n = 3, representative images, scale bar: 10 μm).

### naRNA DAMP activity is dependent on RNA sensors TLR8/Tlr13

To validate naRNA as an immunostimulatory RNA component from the receptor side, we turned to HEK293T cells which do not respond to RNA, unless transfected with plasmids encoding the human ssRNA sensors TLR7 or TLR8 (*20*). NF-κB reporter assays revealed that TLR7 or TLR8-(both ssRNA sensors) but not TLR9 (human DNA sensor)-transfected HEK293T cells stimulated with PMA NETs or mock NETs showed robust NF-κB activity only for PMA NETs (Fig. 3A). R848 (TLR7 and TLR8 agonist) and TL8 or ssRNA+DOTAP (both TLR8 agonists) served as controls. Of note, compared to its cognate agonist, CpG, the DNA sensor TLR9 was only poorly activated by PMA NETs, indicating that naRNA is a superior immune stimulant compared to NET DNA in this system. Indeed, the observed NET response of primary PMNs (Fig. 3B, quantified in C, and Fig. S3A) could be completely blocked by the TLR8-specific inhibitor, CU-CPT9a (*21*), similar to the effects of the non-specific inhibitor, CI-amidine, a PAD4 inhibitor which also blocked the response to PMA (Fig. 3C and S3A). This revealed the RNA sensor TLR8 (*22*) to be the specific naRNA receptor in primary human PMNs. We further genetically validated the involvement of RNA sensing in naRNA-mediated NET propagation using BM-PMNs from *Unc93b1-* or *Tlr13-*defective mice: whereas in *Unc93b1*-defective mice signaling of all endosomal TLRs is abrogated (*23*), mice lacking *Tlr13*, the murine equivalent of TLR8 (*12*), specifically lack ssRNA sensing (*24*). Confocal microscopy analysis showed that WT BM-PMNs responded readily to both PMA (used as control) and PMA NETs, whereas *Unc93b1* (Fig. S3B) or *Tlr13* KO PMNs (Fig. 3D, quantified in E) only responded to the TLR-independent PMA control but not PMA NETs. Additionally, naïve BM-PMNs from *Tlr9* KO mice, which lack DNA sensing, responded to both PMA and PMA NETs with NET formation just like WT BM-PMNs (Fig. S3C), emphasizing the redundancy of DNA vs the key role of naRNA in the self-propagating properties of NETs. This receptor analysis unequivocally confirmed naRNA to be a primary immune stimulant in NETs.

**Fig. 3.**
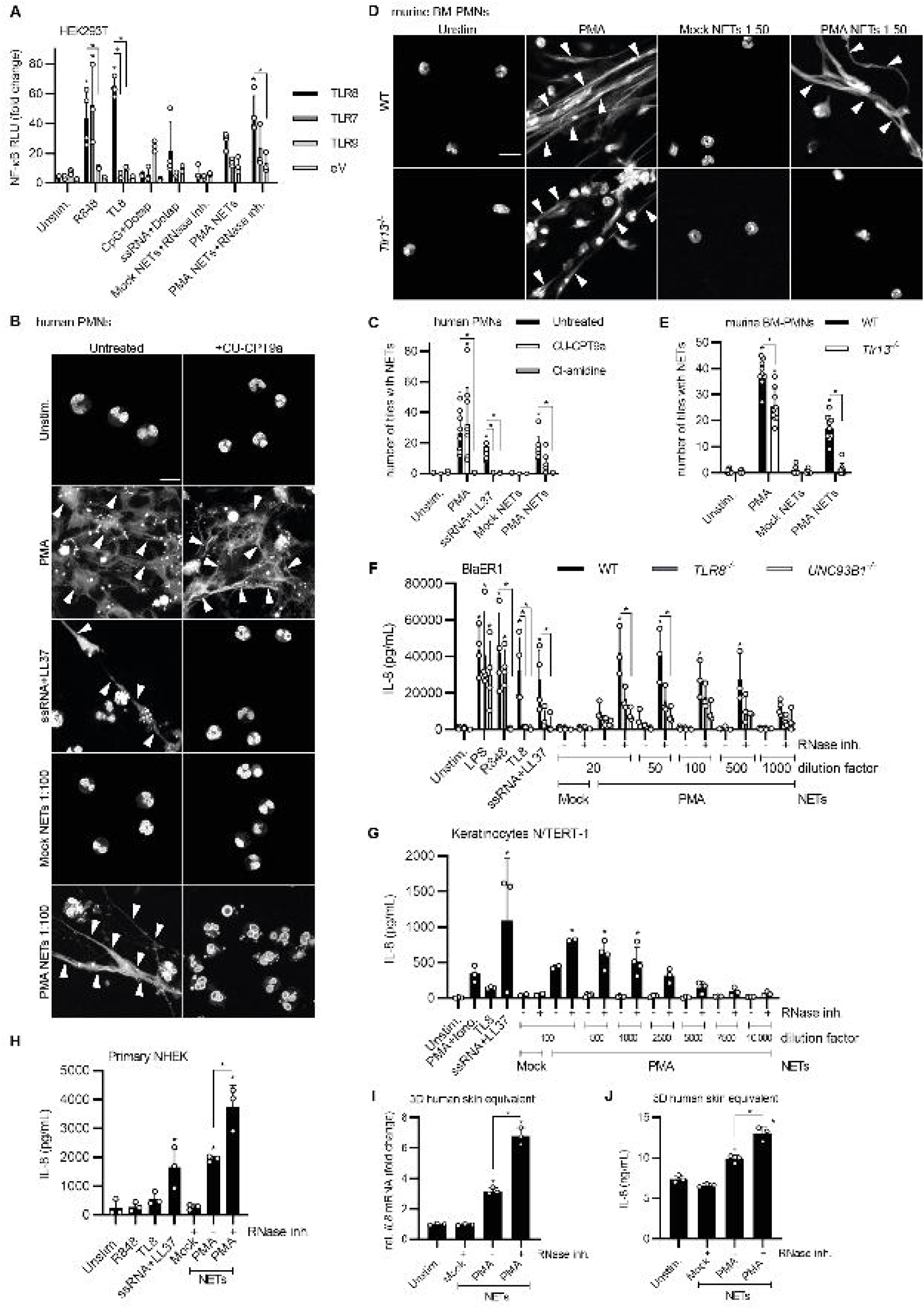
naRNA activity depends on RNA sensors. **(A)** NF-κB dual luciferase assay in HEK293T cell transiently transfected and stimulated as indicated (eV = empty vector, n=3-5, combined data, mean+SD, *p<0.05 according to one-way ANOVA). **(B)** Confocal microscopy of human primary PMNs stimulated as indicated in the presence or absence of CU-CPT9a (100 nM) or CI-amidine (200 μM, not shown) and stained for NETs/DNA (Hoechst 33342, white, n = 3, representative images, scale bar: 10 μm). **(C)** Quantification of **B** using DNA (Hoechst 33342) signal to quantify NET formation (n = 3, combined data, mean+SD, each dot represents the number of NET-positive tiles in one image, three images/condition, *p<0.05 according to non-parametric one-way ANOVA). **(D)** Confocal microscopy analysis of primary C57BL/6 WT or *Tlr13*^*-/-*^ murine BM-PMNs stimulated as indicated and stained for NETs/DNA (Hoechst 33342, white, n = 3, representative images, scale bar: 10 μm). **(E)** Quantification of **D** as in **C** (n = 3, *p<0.05 according to one-way ANOVA). **(F)** Triplicate IL-8 ELISA from WT, *TLR8*^*-/-*^ and *UNC93B1*^*-/-*^ BlaER1 macrophage-like cells stimulated as indicated for 18 h (n = 3-4, combined data, mean+SD, each dot represents one biological replicate, *p<0.05 according to one-way ANOVA). **(G)** Triplicate IL-8 ELISA from N/TERT-1 keratinocytes stimulated as indicated for 24 h (n = 3, combined data, mean+SD, each dot represents one biological replicate, *p<0.05 according to one-way ANOVA). **(H)** As in **G** but with primary normal human epidermal keratinocytes (NHEK) (n = 3, combined data, mean+SD, each dot represents one biological replicate, *p<0.05 according to one-way ANOVA). **(I, J)** Triplicate relative (to unstimulated) *IL8* mRNA qPCR or IL-8 ELISA from NHEK 3D human skin equivalent constructs stimulated as indicated for 24 h (n = 3, representative of one biological replicate is shown, mean+SD, each dot represents one technical replicate, *p<0.05 according to one-way ANOVA).

### PMN-derived naRNA triggers broader immune cell and keratinocyte activation

Given the potent effect of naRNA on naïve PMNs (*cf*. Fig. 3B-E) and the ability of human macrophages to sense RNA via TLR8 (*25*), we hypothesized that naRNA might also directly activate macrophages, which are rapidly recruited at the sites of PMN activation in peripheral tissues (*26*), contributing to an inflammatory process in vivo. We thus assessed the effects of naRNA on genetically modified macrophage-like cell lines: BlaER1 macrophages (*27*) responded to PMA NETs but not mock NETs with IL-8 release, and this was TLR8-dependent as evidenced by comparisons of WT and *TLR8*-edited BlaER1 cells (Fig. 3F). Use of a naRNA-stabilizing RNase inhibitor during NET preparation (*cf*. Fig. 2B) drastically increased the potency of PMA NETs on these macrophages. Additionally, TLR8 and TLR7-edited THP-1 macrophage-like cells (*28*) responded to PMA NETs with lower relative IL-8 release than WT THP-1 cells (Fig. S4A), confirming RNA as the active agent in monocyte/macrophage activation. Human peripheral blood mononuclear cells (PBMCs) were also stimulated with different NET preparations. RNA-stabilized PMA NETs consistently elicited higher TNF, IL-6 and IL-8 levels than non-stabilized PMA- or mock NETs (Fig. S4B-D). However, this effect was not sensitive to the TLR8 inhibitor Cu-CPT9a, which is consistent with the ability of TLR7 to also sense naRNA (*cf*. Fig. 3A) and possibly due to a mixture of cells being present. Further analysis revealed that the natural killer (NK) cell line NK-92 MI showed low IFNγ release in response to RNA-stabilized PMA NETs (Fig. S4E). Collectively, our results show that the RNA component of NETs, naRNA, can engage not only bystander neutrophils in a feed-forward inflammatory response but also other myeloid (monocytes/macrophages) and lymphoid (NK cells) cellular innate immunity. To investigate whether non-hematopoietic cells with immune functions, e.g. tissue cells like keratinocytes, could also respond to RNA-stabilized PMA NETs, N/TERT-1 keratinocytes (*29*) in monolayers were exposed to PMA-NETs (Fig. 3G). This triggered a dose-dependent IL-8 release comparable to synthetic RNA+LL37, especially under RNA stabilization by RNase inhibitors. Similar results were obtained using monolayers of primary normal human epidermal keratinocyte (NHEK) from healthy donors (Fig. 3H). Furthermore, in an ‘in vivo’-like human skin equivalent 3D model in which natural keratinocyte differentiation was recapitulated by NHEK cells (*30*), PMA but not mock NETs significantly induced *IL8* mRNA and protein (Fig. 3I, J). Thus, naRNA can not only activate primary neutrophils in vitro but also trigger broad immune activation in other immune and tissue cells/equivalents.

### naRNA displays DAMP activity in vivo in dependence on RNA sensing

Moreover, to gain an insight into whether naRNA could trigger inflammation in vivo in an RNA receptor-dependent manner, we intradermally injected RNase inhibitor-stabilized mock or PMA NETs in the ears of C57BL6 mice. RNA-stabilized PMA NETs were almost as potent to induce ear swelling as synthetic RNA+LL37 (Fig. 4A), but even non-stabilized PMA NETs induced a response significantly higher than mock NETs. The same stimulants were also injected into the ear of *LysM*^*EGFP/+*^ mice, in which myeloid cells are GFP-positive, thus enabling in vivo monitoring of cellular influx during skin inflammation. Here, RNA-stabilized PMA NETs elicited even greater cellular influx than synthetic RNA+LL37 and mock NETs (Fig. 4B). Importantly, a comparison of WT and *Tlr13*-deficient animals showed that the ear swelling reaction resulting in vivo was naRNA-dependent because RNA-stabilized PMA-NETs were significantly less effective in *Tlr13*-deficient animals (Fig. 4C).

**Fig. 4.**
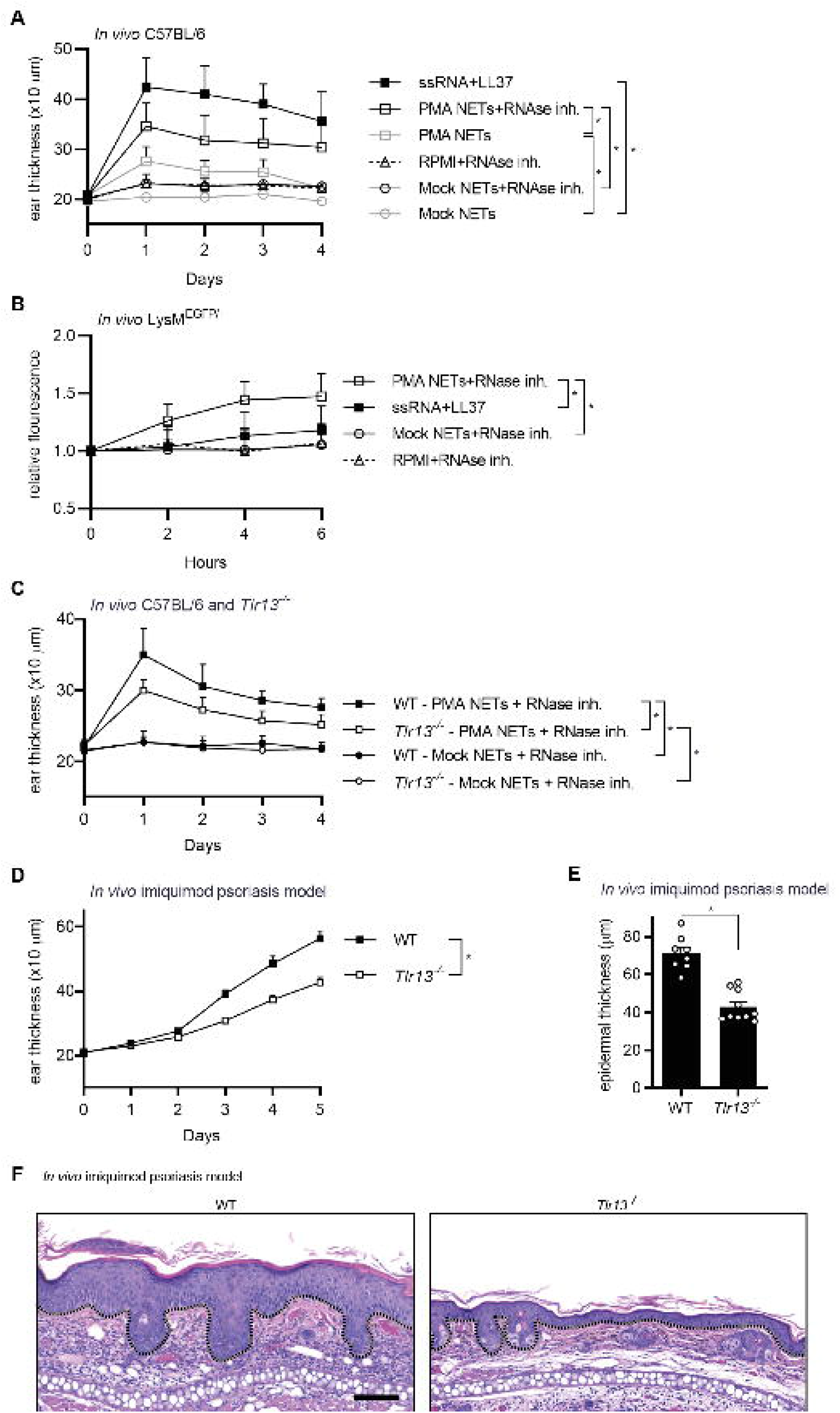
naRNA is a driver of NET-associated *in vivo* inflammation. **(A)** Ear thickness quantified daily in WT C57BL6 mice injected intradermally in vivo on day 0 as indicated (n = 5 per group, combined data, mean+SD, *p<0.05 according to two-way ANOVA). **(B)** Fluorescence imaging monitored hourly in *LysM*^*EGFP/+*^ mice injected intradermally in vivo at t=0 as indicated (n = 10 per group, combined data, mean+SD, *p<0.05 according to two-way ANOVA). **(C)** As in **A** but also using *Tlr13*^*-/-*^ mice (n = 7, combined data, mean+SD, *p<0.05 according to two-way ANOVA). **(D)** As in **C** but instead of intradermal injection with topical imiquimod application on day 0– 4 (C57BL/6 n = 8, *Tlr13*^*-/-*^ n = 10, combined data, mean+SD, *p<0.05 according to two-way ANOVA). **(E)** Measurement of epidermal thickness of H&E-stained ear skin specimens of **D** (C57BL/6 n = 8, *Tlr13*^*-/-*^ n = 10, combined data, mean+SD, *p<0.05 according to Student’s *t*-test). **(F)** Histological analysis of H&E-stained ear skin specimens of **E** (C57BL/6 n = 8, *Tlr13*^*-/-*^ n = 10, representative of one biological replicate is shown, scale bar: 100 μm).

### RNA recognition contributes to progressive skin inflammation in experimental psoriasis

Finally, we sought to explore if naRNA was relevant in an animal disease model. The most frequently used murine model of psoriasis uses the TLR7 ligand imiquimod (IMQ) to trigger increasing skin inflammation characterized by epidermal thickening and immune cell infiltration (*31*). We previously showed that IMQ treatment also leads to the occurrence of NETs in the tissue (*8*). We therefore speculated that naRNA might be involved in disease progression, a hypothesis that could be probed by genetically ablating naRNA sensing and comparing WT and *Tlr13* KO mice. Although in the early induction phase, WT and *Tlr13* KO animals showed a similar increase in ear thickness, the groups diverged from day 3 onwards, after which *Tlr13*-deficient animals were significantly protected from skin inflammation compared to WT mice (Fig. 4D). Additionally, the characteristic epidermal thickening upon IMQ treatment was significantly lower in *Tlr13*-deficient mice compared to WT mice (Fig. 4E, representative image Fig. 4F). The most plausible explanation is that IMQ-initiated NET formation (*8*) amplifies skin inflammation in WT animals via naRNA, whereas this is prevented in *Tlr13* KO animals. This shows that naRNA can significantly contribute to inflammation in a well-established disease model and can act as a potent inflammation-amplifying endogenous DAMP, e.g. in psoriasis.

### naRNA and LL37 await externalization in an already pre-associated DAMP state in healthy donor neutrophils

Thus far, our data aligned well with the notion of the association of LL37 and RNA in the chaotic assembly of the NET accidentally generating a composite DAMP in which LL37 would exert the breaking of tolerance to self-RNA. Indeed, in confocal microscopy (Fig. 5A), Pearson’s colocalization analysis (Fig. 5B) and additional line plot quantification (Figs. 5C and S5) of PMA-stimulated PMN NETs, RNA and LL37 did in fact co-localize in NETs, i.e., after extrusion. However, most surprisingly, in unstimulated PMNs naRNA and LL37 showed even greater Pearson’s colocalization (Fig. 5B) and, unlike DNA, shared co-localization maxima (Fig. 5C, Fig. S5), suggesting the intriguing notion of pre-association of RNA and LL37 even before extrusion. To verify this further, rRNA and LL37 double staining on ultrathin (50-60 nm) sections of resting PMN confirmed the reported localization of LL37 to neutrophil granules (*32*), but also the unexpected concomitant presence of rRNA in the same structures (Fig. 5D). Transmission electron microscopy combined with double staining of rRNA and LL37 using two different immunogold sizes corroborated this further at even higher magnification (Fig. 5E).The concept of pre-packaging of RNA with LL37 in specific compartments also fits with the RNAseq data showing that certain RNA types are lost upon release (*cf*. Fig. 1H), probably because they are not bound to LL37. Of note, the effect was seen in normal peripheral blood PMN across multiple healthy donors, indicating that LL37 association with RNA is a normal physiological PMN property. In turn, and in line with our initial analysis (*cf*. Fig. 1A/C), this would mean that every NET is intended to contain and externalize naRNA-LL37 DAMPs.

**Fig. 5.**
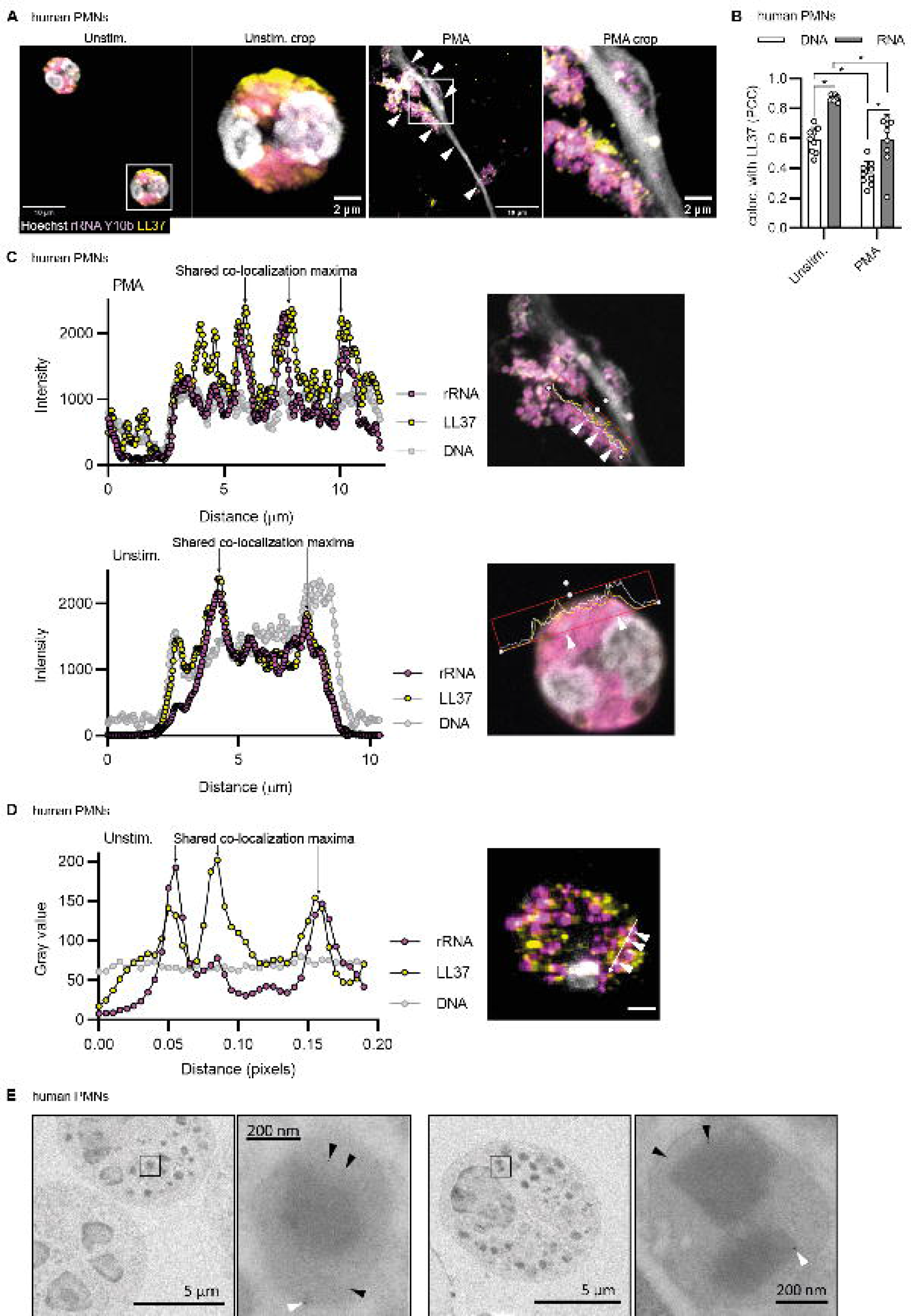
naRNA and LL37 are pre-associated in resting neutrophils. **(A)** Confocal microscopy of primary human PMNs stimulated as indicated for 3 h and stained for naRNA (anti-rRNA Y10b, magenta), LL37 (anti-hLL37-DyLight550, yellow), and DNA (Hoechst 33342, white, n = 3, representative images, scale bar 10 μm). **(B)** Pearson’s correlation coefficient (colocalization) analysis of **A** (n = 3, combined data, mean+SD, each dot represents one image, three images/condition, *p<0.05 according to one-way ANOVA). **(C)** Line plot analysis of LL37, RNA and DNA staining of primary human PMNs stimulated as indicated of **A** was performed using ZenBlue3 software (n=3, representative graph). **(D)** As in **A**/C but on ultrathin sections of unstimulated PMNs (n=3, representative image, scale bar 2 µm). Line plot analysis was performed using ImageJ-Win64 software (n=3, representative graph). **(E)** Transmission electron microscopy of unstimulated human primary PMNs using anti-rRNA and anti-hLL-37 primary and immunogold (6 nm (black arrow) and 12 nm (white arrow) respectively)-labeled secondary antibodies (n = 3, representative images, scale bars as indicated).

## Discussion

At first sight, it may not seem surprising that the release of NETs, a process that churns up the most critical compartment of a cell, e.g., the nucleus, also inevitably leads to the release of another primary cellular biomolecule, namely RNA. However, independently of whether the process is regulated or not, the release of RNA by NETting PMNs appears to have physiological importance as we have demonstrated here. Rather than acting directly in antimicrobial defense (*cf*. Fig. S2A), naRNA appears to be an LL37-associated DAMP of PMN origin that can be released in the NETting process, and then activates both PMN and other immune and tissue cells in an RNA sensor-dependent manner (Fig. S6). Several further findings warrant further discussion:

Firstly, our data revise previous concepts of self-RNA and LL37 in the ‘breaking’ of innate immune tolerance in psoriasis and the physiological role of NETs in general. naRNA when complexed with LL37, gains stimulatory properties, in line with the existing notion that LL37 may ‘break’ immune tolerance by acting as a physiological ‘transfection reagent’ for uptake into immune cells, and by shielding it from RNase degradation. However, our data show that RNA and LL37 do aggregate and co-localize ‘accidentally’ in the chaotic mesh of the NET: quite unexpectedly, they were rather found in a kind of ‘pre-association’ in the same intracellular storage compartments before extrusion during NET formation. It will be interesting to study the process for pre-association for this first ‘composite DAMP’ and elucidate the precise nature of the RNAs assembled with LL37 in the future. But already the identification of such pre-association challenges existing concepts of LL37-RNA complexes as ‘accidentally assembled’ tolerance breakers in certain diseases (*10*). Rather, it suggests naRNA-LL37 complexes act as purposefully pre-associated DAMPs that arm neutrophils at steady state and are meant to ‘break tolerance’ and cause immune-activation in healthy donors – with other pathomechanism causing chronic disease in e.g. psoriasis patients. Especially since the here-described phenomena were found with neutrophils and other cells from healthy donors, intentional (rather than accidental/extracellular) pre-association points to a very specific but broadly relevant novel role of the naRNA-LL37 DAMP in NET biology: The fact that NETs have been described for a multitude of sterile conditions, would anyways argue that antimicrobial cannot represent the primary, let alone only, intended purpose of this widespread phenomenon. Rather, based our findings, we believe the NET response in general to primarily be a DAMP response, calling for a re-evaluation of existing concepts in NET biology. We speculate that the physiological relevance of deploying this composite DAMP upon NET formation could be to tag fresh NETs with a timed molecular beacon: In this scenario, freshly extruded NETs decorated with the naRNA-LL37 DAMP would highlight an acute tissue insult (or the lingering presence of a pathogenic microorganism during non-sterile conditions) to other cells not directly or initially engaged by the threat. The naRNA-LL37 DAMP would thus label ‘fresh NETs’ as ‘requiring attention’ and trigger further immune activation, clearance and eventually inflammation resolving activities. Over time, whilst the DNA-related structural and antimicrobial properties of NETs may remain longer, inevitable RNA degradation would lead to deactivation of the composite DAMP, rendering ‘old NETs’ less immunostimulatory. Further work outside the scope of the present study may explore the intriguing notion of naRNA as time-wise self-restricting molecular label for NETs.

Moreover, our data pertain to the origin and effects of ‘extracellular RNA’ (exRNA). exRNA is a generic term to indicate a heterogeneous group of RNA molecules which are actively or passively released during sterile inflammation or infectious processes. exRNA can be released in a ‘free’ state, bound to proteins or phospholipids, in association with extracellular vesicles (EVs) or apoptotic bodies (*14*). In all these forms, exRNA may function as a DAMP but also as e.g., procoagulant or regenerative factor (*14*). Our data identify naRNA as the first type of exRNA for which a clearly defined physiological origin/process (namely NETs) of extracellular release in physiologically relevant amounts is provided. We speculate that our findings will help trace multiple descriptions of exRNA, on the one side, to neutrophil traps on the other. For example, exRNA has emerged as disease-relevant in atherosclerosis, where it was described to act as a proinflammatory mediator enhancing the recruitment of leukocytes to the site of atherosclerotic lesions as shown in a mouse model of accelerated atherosclerosis (*33*). At the same time, NETs have been ascribed a role in amplifying sterile inflammatory responses in an independent mouse model of atherosclerosis by priming macrophages (*3*). However, never have these two independent strains of research been connected. By showing NETs to release naRNA, a DAMP type of exRNA, our work connects both lines of enquiry. Likewise, for rheumatoid arthritis (RA), our work makes a plausible link between exRNA in synovial fluid contributing to joint inflammation (*34*), the emerging role of NETs in RA (*4*) and even a hitherto enigmatic but therapeutically relevant role of TLR8 (*35, 36*).

Moreover, NETs have been of broad interest focus in COVID-19-related research regarding thrombo-inflammatory states, like sepsis, thrombosis, and respiratory failure,, have been found in plasma of hospitalized COVID-19 patients, and linked to thrombus formation by several research groups (*37*) (*38*) (*39*) (*5, 40*). Furthermore, exRNAs have been in focus as procoagulant cofactors in blood coagulation (*41*) (*42*). A link between NET-related and RNA-induced thrombus formation in COVID-19 has never been made but highlights naRNA as a potential contributing factor of NET-induced thrombo-inflammatory states in COVID-19 disease pathogenesis. Collectively, we suspect similar links between NETs and exRNA via naRNA to emerge for cardiovascular diseases and cancer if the role of naRNA is thoroughly assessed. Furthermore, the broad sensitivity of immune and tissue cells to naRNA observed by us makes sense of how exRNA may act pathophysiologically.

Although we believe PMNs to be the primary trap forming human leukocyte population and hence naRNA the most significant “trap-associated RNA”, it will be interesting to explore whether mast cell (MCETs) (*43*) or macrophage extracellular traps (METs) (*44*) contain RNA. However, unlike PMN, the latter immune cells are not primary sources of LL37 (*32*), so that RNA associated with MCETs or METs would be of lesser physiological relevance as a DAMP than naRNA due to a lack of LL37. Therefore, translational approaches, e.g., to restrict trap-RNA mediated amplification of inflammation, should probably center on PMN-derived naRNA. The use of PAD4 inhibitors has already been investigated in animal models to treat cancer or atherosclerotic lesions (*45, 46*) and would represent one potential way of eliminating NETs and hence naRNA-mediated inflammation. However, this would also prevent the effects of NETs that are beneficial for host defense (e.g. physical trapping via DNA and antimicrobial enzymatic activities) and may render treated patients vulnerable to infections. From a translational perspective, our in vivo data indicate that blockade of RNA sensing might be more advantageous, restricting only naRNA-mediated responses. Although probably not applicable to patients, it is underscored by evidence that neuroinflammation upon subarachnoid hemorrhage, which is characterized by a NET pathology, is strongly ameliorated by RNase treatment in vivo (*47*). More applicable to patients may be the targeting of TLR8 by small molecular antagonists (*48*) or inhibitory oligonucleotides, which are able to block PMN responsiveness to synthetic RNA in vitro (*8*). Exciting is thus the observed efficacy of TLR inhibitory oligoribonucleotides in a psoriasis mouse model (*49*) and even a first clinical trial in psoriasis patients (*50*). We could imagine the blockade of naRNA effects via TLRs highlighted here to emerge as an effective novel strategy to target multiple exRNA or NET-related inflammatory responses.

## Aditional information

### Author contributions

according to Credit guidelines:

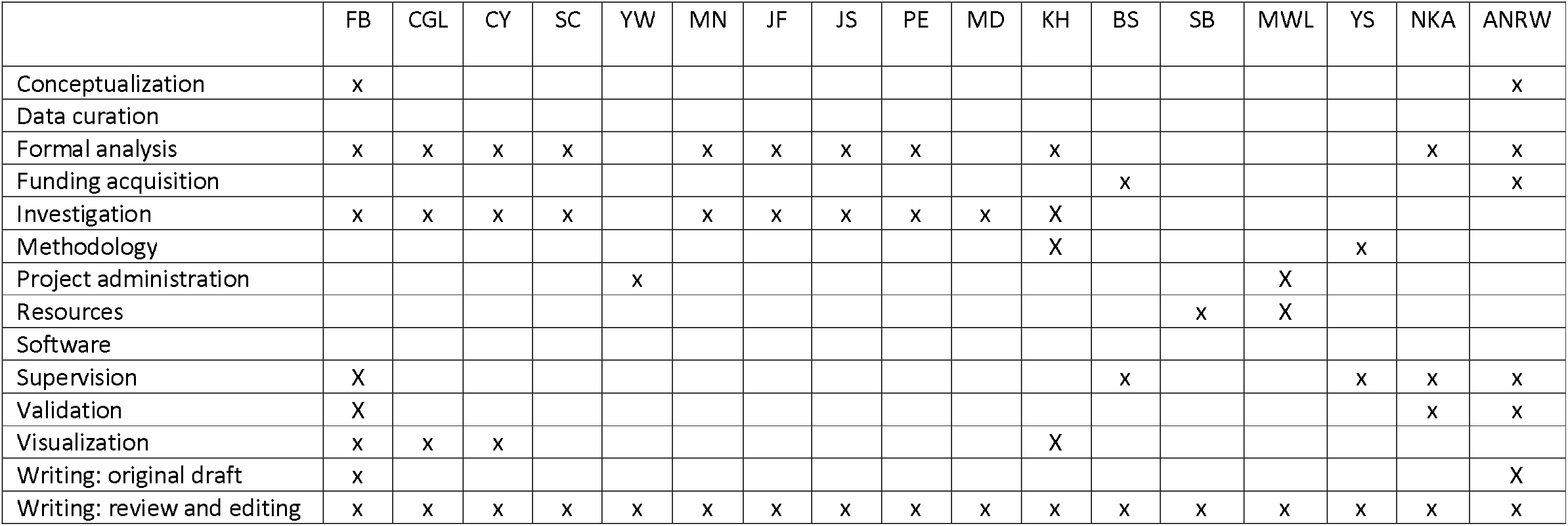

## Conflict of interest statement

NKA has received previous grant support from Pfizer and Boehringer Ingelheim and was a paid consultant for Janssen Pharmaceuticals. All other authors declare no competing interests. S.D.G.

Acknowledgments

We gratefully acknowledge Jim Rheinwald, Holger Heine, Austin Chang, Thomas Zillinger for the provision of reagents, respectively, and Jon Kagan, Alexander Dalpke and Libera Lo Presti for helpful scientific and editorial comments. We further acknowledge Bettina Danker, Mark Helm and Martina Christina Schmidt-Dengler for very helpful technical and scientific support. We thank all voluntary healthy donors of biomaterials for participating in the study. The study was supported by the Deutsche Forschungsgemeinschaft (German Research Foundation, DFG) grants CRC TR156 “The skin as an immune sensor and effector organ – Orchestrating local and systemic immunity” (to FB, CG, JF, JS, BS and ANRW), NIH grants R01AI146177, R01AR073665, and R01AR069502 (to NKA). NKA. has received previous grant support from Pfizer and Boehringer Ingelheim and was a paid consultant for Janssen Pharmaceuticals. Infrastructural funding was provided by the University of Tübingen, the University Hospital Tübingen and the DFG Clusters of Excellence “iFIT – Image-Guided and Functionally Instructed Tumor Therapies” (EXC 2180, to AW, PE, BS and MWL), “CMFI – Controlling Microbes to Fight Infection (EXC 2124 to AW and BS). Gefördert durch die Deutsche Forschungsgemeinschaft (DFG) im Rahmen der Exzellenzstrategie des Bundes und der Länder - EXC 2180 and EXC 2124. The authors declare no competing financial interests.

### Abbreviations

AF: Alexa Fluor;
bRNA: bacterial ribonucleic acid;
DAMP: damage-associated molecular pattern;
5- EU: 5-Ethynyluridine;
FACS: Fluorescence Activated Cell Sorting;
fRNA: fungal ribonucleic acid;
H&E: Hematoxylin and Eosin;
HEK: Human embryonic kidney;
HMGB-1: High-Mobility-Group-Protein B1;
HSC: hematopoietic stem cell;
IFN: Interferon;
IL: Interleukin;
KO: knockout;
MCET: Mast cell extracellular trap;
MET: Macrophage extracellular trap;
MPO: Myeloperoxidase;
naRNA: Neutrophil extracellular trap-associated RNA;
NET: Neutrophil extracellular trap;
NK: natural killer;
PAD4: Peptidyl arginine deiminase 4;
PBMC: Peripheral Blood Mononuclear Cell;
PKC: Protein kinase C;
PMN: Polymorphonuclear leukocytes;
RT: room temperature;
SEM: scanning electron microscopy;
TLR: Toll-like receptor;
TNF: Tumor necrosis factor

## Full online materials and methods

### Reagents

PMA (tlrl-pma), Nigericin (tlrl-nig), LL37 (tlrl-l37), as well as the PRR ligands LPS (tlrl-peklps), R848 (tlrl-r848), TL8 (tlrl-tl8506), and the TLR8-inhibitor CU-CPT9a (inh-cc9a) were from Invivogen, Ionomycin was acquired from Sigma (I0634-1MG). RNase inhibitor (N2615) was from Promega, RNase A (EN0531), DNase I (EN0521) and DNase inhibitor (EN0521) were from Thermo Fisher. The PAD4-inhibitor Cl-amidine (506282) was from Merck Millipore. DOTAP (L787.2) was from Roth (see Supplementary Table S1), ssRNA40 was from Eurogentec and CpG PTO 2006 from TIB Molbiol (see Supplementary Table S2). Bacterial RNA isolated from *S. aureus* was prepared as described (*8*). Fungal RNA from *C. albicans* strain SC5314 was isolated as described below, as well as naRNA isolated from PMA NETs. For complex formation to stimulate cells in a volume of 500 μL, 5.8 μM ssRNA40 (∼ 34.4 μg/mL), fungal RNA (125 ng/mL), bacterial RNA (10 μg/mL) or PMA NET derived naRNA (∼ 600 ng/mL) was mixed with 10 μg LL37 and left for 1 h at room temperature (RT). For a smaller volume of cells, complexes were used in the according fractional amount. For RNA-only or LL37-only controls, the same amounts and volumes were used replacing one of the components with sterile, endotoxin-free H_2_O. For complex formation with DOTAP, the according RNA or CpG was incubated with the transfection reagent for 10 min at RT prior to stimulation of the cells. NET content for stimulation was prepared as described below. Antibodies used for fluorescence microscopy, as well as click chemistry reagents are listed in Supplementary Tables S3 and S4. Constructs used for transfection of HEK293T cells are listed in Supplementary Table S5.

### Preparation of fungal RNA from *C. albicans*

*C. albicans* SC5314 (kindly cultured and prepared by Tzu-Hsuan Chang, Tübingen) was plated in a slant tube containing YPD agar and grown overnight at 30 °C as described in (*51*). One colony was picked from the slant tube and resuspended in 500 μL YPD medium, centrifuged at 10,000 rpm for 1 min and washed with phosphate-buffered saline (PBS). Afterwards, the pellet was resuspended in 200 μL RLT buffer (derived from RNeasy Mini Kit, Qiagen, #74106) and transferred into a 2 mL tube containing 0.5 mm diameter ceramic beads. The tube was filled up to 1 mL with RLT buffer and the fungi were subsequently homogenized by using a microtube homogenizer (BeadBug™, Merck) with an interval of seven times 2 min shaking at 2800 rpm and 1 min cooling break on ice. The supernatant was transferred into a new tube containing 1 mL 75% ethanol (VWR, 20821.330). The further RNA isolation was performed according to the manufacturer’s instructions using the Qiagen RNeasy Mini Kit for purification of Total RNA from Animal Tissues (RNeasy Mini Kit, Qiagen, 74106). The RNA was eluted in 30 μL RNase DNase-free H_2_O and the concentration was determined with a Nanodrop Spectrophotometer.

### Mice

*Unc93b1*^*3d/3d*^ (*23*), *Tlr13*^*-/-*^ (*24*) (both kindly provided by Tatjana Eigenbrod, Heidelberg and on C57BL/6 background) and WT C57BL/6 mice between 8 and 20 weeks of age were used in accordance with local institutional guidelines on animal experiments, regular hygiene monitoring, and specific locally approved protocols compliant with the German regulations of the Gesellschaft für Versuchstierkunde/Society for Laboratory Animal Science (GV-SOLAS) and the European Health Law of the Federation of Laboratory Animal Science Associations (FELASA) for sacrificing and *in vivo* work. *Unc93b1*^*3d/3d*^, *Tlr13*^*-/-*^ and WT C57BL/6 control mice were housed in controlled specific-pathogen-free animal facilities at the Interfaculty Institute of Cell Biology, Tübingen. Local federal authority for the approval of experimental protocols was the Regierungspräsidium Tübingen. *Tlr9*^*-/-*^ mice and matched WT C57BL/6 control mice were a kind gift from Birgit Schittek, Tübingen. LysM^EGFP/+ 52^(*52*)(*52*), *Tlr13*^-/-^ mice (*24*) (kindly provided by James Chen, Houston and David Nemazee, La Jolla) and WT (all on a C57BL/6 background) mouse strains were bred and maintained under the specific pathogen-free conditions, with air isolated cages at an American Association for the Accreditation of Laboratory Animal Care (AAALAC)-accredited animal facility at Johns Hopkins University and handled according to procedures described in the Guide for the Care and Use of Laboratory Animals as well as Johns Hopkins University’s policies and procedures as set forth in the Johns Hopkins University Animal Care and Use Training Manual, and all animal experiments were approved by the Johns Hopkins University Animal Care and Use Committee (MO21M378). Gender-and age-matched 6–8-week-old mice were used for each experiment.

### Isolation and stimulation of primary bone-marrow-derived polymorphonuclear neutrophils (BM-PMNs)

Bone-marrow (BM)-PMNs were isolated from the bone marrow as described (*8*). In brief, bones were isolated from the respective mice and the bone marrow was flushed out. Afterwards, neutrophils were isolated using magnetic separation (mouse Neutrophil isolation kit, Miltenyi Biotec, 130-097-658) following the manufacturer’s instructions. In total, 11.×1.10^5^1.cells/well PMNs were seeded in a 24-well plate, and stimulation was carried out with PMA (600 nM), ssRNA+LL37 complex (as previously described), nigericin (50 μM), live *C. albicans* (MOI1) or human NET content (mock control and PMA NETs, 1:50 dilution) for 161.h at 37°C and 5% CO2. Subsequently, cells were stained for immunofluorescence.

### Study participants and human blood or tissue sample acquisition

All healthy donors included in this study provided their written informed consent before participation. Approval for use of biomaterials was obtained for this project by the local ethics committee of the Medical Faculty Tübingen in accordance with the principles laid down in the Declaration of Helsinki as well as applicable laws and regulations.

### Primary human neutrophil isolation and stimulation

Neutrophils of healthy human donors were isolated as described (*8*). In brief, EDTA-anticoagulated whole blood was diluted in PBS (Thermo Fisher, 14190-169), loaded on Ficoll (1.077 g/mL, Sigma, 10771) and centrifuged for 25 min at 509 x g at RT without brake. Afterwards, all layers except the erythrocyte-granulocyte layer were discarded and erythrocyte lysis was performed twice (for 20 and for 10 min) using 1x ammonium chloride erythrocyte lysis buffer (see Supplementary Table S6) at 4 °C on roller shaker. The remaining cells were resuspended in culture medium (RPMI culture medium (Sigma-Aldrich, R8758)⍰.+ ⍰.10% FBS (heat-inactivated, sterile filtered, Th. Geyer, 11682258)) to a concentration of 1.6 × 10^6^ cells/mL. 500 μL of cells were seeded in a 24-well plate for immunofluorescence microscopy or 8 mL of 5 × 10^6^ cells/mL in 10 cm uncoated dishes for NET preparation and naRNA isolation/isolation of whole PMN RNA. After seeding, the cells were rested for 30 min at 37°C and 5% CO_2_, followed by 3 h stimulation with PMA (600 nM), nigericin (50 μM), live *C. albicans* (MOI2), RNA+LL37 complexes (as previously described) or NET content at indicated dilutions for IF or 4 h stimulation with PMA (600 nM) for NET preparation. Where indicated, the cells were incubated with 100 nM TLR8-inhibitor CU-CPT9a or 200 μM PAD4-inhibitor Cl-amidine 30 min before stimulation and medium was not replaced during incubation with the respective stimuli.

### Primary peripheral blood mononuclear cell (PBMC) isolation and stimulation

PBMCs were isolated from whole blood or buffy coats as described (*8*). In brief, EDTA-anticoagulated blood was diluted in PBS and density gradient separation was performed as described above. The PBMC layer was then carefully transferred into another reaction tube and diluted in PBS (1:1). The cell suspension was spun down at 645⍰.× ⍰.g for 8⍰.min and the cells were washed two more times in PBS and resuspended in culture medium (RPMI⍰.+ ⍰10% FBS (heat inactivated) +1% L-glutamine) at a density of 1 × 10^6^ cells/mL. Afterwards, 200 μL of PBMCs were seeded in a 96-well plate and stimulated with LPS (100 ng/mL), R848 (5 μg/mL), TL8 (100 ng/mL), ssRNA (1.6 μg/mL) + DOTAP (50 μg/mL), ssRNA+LL37 complex (as described above), and respective NET content (1:20 dilution) for 24 h at 37°C and 5% CO_2_. Where indicated, the TLR8 inhibitor CU-CPT9a was added to the cells at a concentration of 1 μM 2 h before stimulation and was not removed for the incubation of the cells with the respective stimuli. After the stimulation, the plate was centrifuged for 5 min at 1500 rpm and the supernatant was collected and stored at -80 °C until the ELISA was performed.

### Preparation of NETs and isolation of naRNA/whole PMN RNA

NETs were prepared by 600 nM PMA treatment for 4 h at 37°C and 5% CO_2_, or cells were left untreated during the incubation as the mock control. After incubation, the neutrophils were gently washed three times with PBS to get rid of PMA as the stimulus for NET formation, any cytokines released by the cells and unstimulated PMNs, as those do not adhere to the uncoated petri dish used here. In some conditions (as indicated) during the NET preparation process and for storage, naRNA was protected by addition of 10 U/μL RNase inhibitor (= mock or PMA NETs + RNase Inhibitor). For digest of NETs, the NET content was incubated for 20 min at 37 °C with RNase A (Thermo Fisher, EN0531; 100 μg/mL, EDTA for DNase inhibition added) at 37 °C. For isolation of naRNA, RNase inhibitor was added to the PMNs during NET formation. After the above-described washing process, PMA or mock NETs were resuspended in 300 μL of ML buffer and RNA isolation was performed according to the manufacturer’s instructions (NucleoSpin miRNA isolation kit, Macherey-Nagel, 740971.50). The RNA was eluted in 50 μL RNase DNase-free H_2_O and the concentration was determined with a Nanodrop Spectrophotometer. For isolation of whole PMN RNA, untreated PMNs were directly lysed, and the RNA was isolated from the cells according to the manufacturer’s instructions.

### Preparation of human primary stem cell-derived PMNs

Stem cells derived from human healthy donors were prepared and differentiated as described (*19*). In brief, bone marrow was diluted in PBS, carefully layered on Ficoll-Paque medium (density: 1.077 g/mL) and centrifuged at 500 x g for 25 min at RT without brakes. The interphase layer containing the mononuclear cell fraction was transferred to a new tube and washed twice with 30 mL ice-cold PBS by centrifugation at 300 x g for 8 min at 8 °C. Further, the cells were resuspended, counted, and isolated using Human CD34 MicroBead Kit (Miltenyi) for magnetic beads-based isolation of CD34^+^ cells from BM-MNCs. Afterwards, the number of CD34^+^ HPSCs was determined and cultured in CD34^+^ culture medium (Stemline II Hematopoietic Stem Cell Expansion medium supplemented with 10% FCS, 1% L-glutamine, 1% penicillin/streptomycin, and a human recombinant cytokine cocktail consisting of 20 ng/mL IL-3, 20 ng/mL IL-6, 20 ng/mL TPO, 50 ng/mL SCF, and 50 ng/mL FLT-3L) at a density of 2 × 10^5^/mL at 37 °C and 5% CO_2_. The medium was replaced every second day and the cells were cultured for 14 days. During the differentiation process, the cells were treated with 100 μM 5-ethynyluridine for 14 days for subsequent click chemistry labeling of endogenous RNA or were left untreated as negative controls. To verify differentiation, cell morphology was assessed using Cytospin assay. In brief, the Cytoclip™ slide clips were loaded by fitting the filter card, the sample chamber, and the glass slide. An assemble Cytoclip™ slide clip was then placed in the slide clip support plate of the cytospin centrifuge. 2 × 10^4^ cells from liquid culture differentiation were pipetted into Cytofunnel™ and centrifuged for 3 min at 200 x g. The cytospin slides were stained for 5 min in May-Grünwald stain, rinsed shortly with ddH_2_O, and then stained for 10 min in Wright-Giemsa stain. Afterwards, the slides were rinsed shortly with ddH_2_O, and cell morphology was determined using a microscope. To further verify differentiation, flow cytometric analysis was performed using antibodies specific for the following hematopoietic/myeloid markers: CD45 (leukocyte marker), CD34 (HSPC marker), CD33 (promyelocyte marker), CD11b (myeloid cell marker), CD14 (monocyte marker), and CD15 and CD16 (neutrophil markers). Neutrophil percentage was determined by gating on neutrophils as follows: CD45^+^CD11b^+^CD15^+^, or CD45^+^CD11b^+^CD16^+^, or CD45^+^CD15^+^CD16^+^ cells

### BlaER1 cell culture, transdifferentiation and stimulation

BlaER1 cells (WT, *Unc93b*^*-/-*^ and *Tlr8*^*-/-*^), a kind gift of Holger Heine, Borstel, Germany (*27*), were cultured, transdifferentiated, and stimulated for 18 h with the respective stimuli as described (*8*). In brief, 1 × 10^6^ cells/well were seeded in a 6-well plate and differentiated with 10 ng/mL hIL-3, 10 ng/mL hMCSF and 150 nM β-estradiol in complete RPMI-1640 (PANBiotech, P04-18525) for 7 days. Afterwards 5 × 10^4^ differentiated cells were reseeded in a 96-well plate, followed by 1 h resting. Cells were treated with the respective stimuli (LPS at 0.1 μg/mL, R848 at 5 μg/mL, TL8 at 100 ng/mL or ssRNA+LL37 complex as described above) and mock or PMA NETs with or without RNase inhibitor) in complete medium in a total volume of 125 μL/well for 18 h. After stimulation, the cells were centrifuged for 5 min at 1200 rpm, the supernatant was transferred into a new plate and stored at - 80°C until the ELISA was performed.

### THP-1 cell culture, differentiation and stimulation

THP-1 cells (THP-1 cells were a kind gift from Thomas Zillinger, Bonn, Germany (*28*)), were cultured in complete RPMI-1640 (Sigma, R8758-24X500ML) medium. For differentiation into macrophage-like cells, 5 × 10^4^ cells/well were seeded in a 96-well plate, treated with 300 ng/mL PMA and incubated for 16 h at 37°C and 5% CO_2_. The next day, the cells were washed three times with PBS and fresh medium was added, followed by 48 h of resting. Subsequently, the medium was removed, exchanged by medium containing 200 U/mL IFN-γ (Sigma, I-3265), and the cells were incubated for 6 h. After repeated washing and medium exchange, cells were treated with the respective stimuli (PMA (25 μg/mL) + Ionomycin (0.375 μg/mL), LPS (0.1 μg/mL), R848 (5 μg/mL), TL8 (40 ng/mL), ssRNA+LL37 complex (as described above), mock NETs + RNase inhibitor (1:50 dilution), PMA NETs (1:50 dilution) and PMA NETs + RNase inhibitor (1:50 dilution)) in complete medium in a total volume of 125 μL/well for 18 h. After stimulation, the cells were centrifuged for 5 min at 1200 rpm, the supernatant was transferred into a new plate and stored at -80°C until the ELISA was performed.

### N/TERT-1 keratinocyte cell culture and stimulation

N/TERT-1 cells (a kind gift from Prof. James Rheinwald (*29*)) were cultured for less than ten passages in complete CnT-07 medium (CELLnTEC, CnT-07). Two days prior to stimulation, a total amount of 2 × 10^4^ cells/well was seeded in a 96-well plate and incubated at 37°C and 5% CO_2_. The medium was renewed, and the cells were stimulated in a total volume of 125 μL/well of the respective stimuli diluted in medium (PMA (25 μg/mL) + Ionomycin (0.375 μg/mL), TL8 (200 ng/mL), ssRNA+LL37 complex (as previously described), as well as mock and PMA NETs with and without RNase inhibitor at indicated dilutions). After 24 h stimulation, the cells were centrifuged at 1200 rpm for 5 min and the supernatant was stored in a new plate at -80°C until further usage.

### Primary normal human epidermal keratinocyte (NHEK) cell culture and stimulation

NHEK cells (Normal Human Epidermal Keratinocytes (NHEK) single juvenile donor, proliferating, PromoCell, C-12002) were grown and stimulated in Keratinocyte Growth Medium 2 (PromoCell, C-20111). Two days prior to stimulation, a total amount of 2 × 10^4^ cells/well was seeded in a 96-well plate and incubated at 37°C and 5% CO_2_. For stimulation, the medium was renewed with basal medium (PromoCell, C-20211) containing 1.7 mM CaCl_2_ (Roth, CN93.1) and stimuli were added and diluted in medium in a final volume of 125 μL (R848 (20 μg/mL), TL8 (200 ng/mL), ssRNA+LL37 complex, as well as mock NETs + RNase inhibitor (1:25 dilution) and PMA NETs with and without RNase inhibitor (1:25 dilution)). After 24 h stimulation, the cells were centrifuged at 1200 rpm for 5 min and the supernatant was stored in a new plate at -80°C until further usage.

### Preparation of 3D human skin equivalent

The 3D human skin equivalent was prepared as described (*53*). Briefly, primary fibroblasts were seeded on collagen and incubated in FF medium for five days. Subsequently, primary keratinocytes were added to the wells and airlifting was performed on day 12. On day 22, the 3D skin model was stimulated with NET content (25 μL/well for one 3D construct grown in a 12-well chamber) or respective water control for 24 h. Afterwards, supernatant was harvested for ELISA, RNA was isolated for qPCR analysis and H&E staining was performed.

### NK-92 MI cell culture and stimulation

NK-92 MI cells (kindly provided by Melanie Märklin, University Hospital Tübingen) were cultured in IMDM-Medium (Lonza, 12-722F). For stimulation, a total amount of 1 × 10^5^ cells/well were seeded in a volume of 200 μL and rested for 2 h at 37°C and 5% CO_2_. The cells were stimulated with LPS (100 ng/mL), CpG (2.5 μM) + DOTAP (25 μg/mL), R848 (5 μg/mL), TL8 (100 ng/mL), ssRNA (1.6 μg/mL) + DOTAP (50 μg/mL), ssRNA+LL37 complex (as described above), or NET content (1:100 dilution) for 24 h and afterwards centrifuged for 5 min at 1500 rpm. Subsequently, the supernatant was transferred into a new plate and stored at -80°C until further usage.

### Flow cytometry

After PMN isolation, the purity and activation status of the cells was determined by flow cytometry as described (*8*). In brief, 200 μL of cells were transferred into a 96-well plate (U-bottom) and centrifuged for 5 min at 448 x g. Afterwards, blocking was performed using pooled human serum diluted 1:10 in PBS for 15⍰.min at 4°C. After washing, the samples were stained for 201.min at RT in the dark and fixed (4% PFA in PBS) after repeated washing for 10⍰.min at RT in the dark. After an additional washing step, the cell pellets were resuspended in 300⍰.µL PBS and measurements were performed on a FACS Canto II (BD Bioscience, Diva software). Analysis was performed using FlowJo V10 analysis software.

### RNA sequencing analysis of naRNA

Isolated naRNA was analyzed for quality control using the Agilent 4200 TapeStation system. Subsequently, the RNA was sequenced according to NEBNext® Ultra™ II Directional RNA Library Prep Kit for Illumina® using the protocol for use with rRNA Depleted FFPE RNA. The data was quantified using Salmon Version1.5.0 and tximport was used to obtain the transcript-level quantification. For transcript classification, GENECODE annotation was performed.

### Fluorescence microscopy of fixed human or murine primary neutrophils

500 μL of 1.6 × 10^6^ cells/mL of human blood PMNs, and 2 × 10^6^ cells/mL murine BM-PMNs were seeded in a 24-well plate containing poly-L-lysine-coated glass coverslips (Electron Microscopy Sciences, 72292-04) and rested for 30 min before stimulation at 37°C and 5% CO_2_ with the according stimuli (as described above) for 3 h (human) or 16 h (murine), respectively, as adapted from Brinkmann *et al*., 2004 (*1*). After stimulation, the cells were carefully washed with PBS and fixed with fixation buffer (Biolegend, 420801) for 10 min at RT in the dark. Afterwards, the cells were blocked with PBS containing 0.1% heat-inactivated diethylpyrocarbonate (DEPC) (Roth, K028.2), 10% chicken serum (Normal Chicken Serum Blocking Solution S-3000, Biozol/Vectorlabs, VEC-S-3000-20), 0.1% saponin (Applichem, A4518.0100), as well as 10 U/μL RNase inhibitor for 2 h at RT. The primary antibodies (rRNA Y10b, hLL37, see Supplementary Table S3) were diluted 1:50 in blocking buffer and subsequently incubated for 2 h at RT. Afterwards, the cells were washed three times with PBS containing 0.1% heat inactivated DEPC and incubated with the secondary antibodies (see Supplementary Table S3) in a 1:500 dilution in blocking buffer for 1 h. After repeated washing, the cells were incubated with Hoechst 33342 (Thermo Fisher; 1 μg/mL) for 5 min to stain nuclear DNA. Secondary antibodies alone did not yield any significant staining under identical staining and acquisition conditions. The coverslips were mounted (ProLong™ Diamond Antifade Mountant, Thermo Fisher, P36961) on glass slides and left to dry overnight at RT in the dark. Subsequently, the samples were stored at 4°C before microscopy using a Zeiss LSM800 Confocal microscope (40x or 63x magnification with z-stack acquisition, AiryScan mode) and image analysis using ImageJ-Win64 and Zen Blue3 software was performed. For IF of ultrathin sections, coverslips were coated with a carbon layer, glow-discharged and UV-treated overnight before cells were seeded. After incubation, the cells were fixed in 2.5% glutaraldehyde in PBS for 2 h at room temperature followed by incubation at 4°C. Samples were prepared by the progressive lowering of temperature method (PLT). Therefore, the fixed cells were incubated in ethanol for infiltration of Lowicryl HM20, followed by UV polymerization and subsequent sectioning into 50-60 nm ultrathin slices. Immunolabeling was performed by incubation with anti-rRNA Y10b and anti-LL37 primary antibodies for 1 h at RT. Afterwards, the cells were incubated with goat anti-mouse Cy3 and goat anti-rabbit AF488 in blocking buffer for 1 h at RT. Following immunolabeling, the samples were treated with 1% uranyl acetate for 5 min at RT before analysis using a Zeiss Axioplan microscope (63x objective, Olympus SIS cell software) and image/line plot analysis was performed using ImageJ-Win64.

### Quantification of NET formation

To quantify the formation of NETs, microscopy images were obtained using a Zeiss LSM800 Confocal microscope with a 40x objective and 3x3 tiles acquisition. Three images per sample of three biological replicates were taken. To quantify NET formation by using NET-related signal dispersion of rRNA and DNA signal, ImageJ-Win64 software was used, and a threshold (Triangle threshold) was applied as originally described (*54*). Particles were analyzed with a ROI (region of interest) manager (size (micron^2^): 100-infinity (pixel units); circularity 0.00-1.00) and average size and number of particles (ROIs) were assessed. In NETs, RNA and DNA signals showed up in a greater number and smaller size, making the usage of the ratio suitable as a measurement of NET-related signal dispersion. In the case of using the DNA signal only to assess NET formation, ImageJ-Win64 software was used to create a PNG image and a grid with 8x8 tiles was manually applied to the images. Tiles containing extracellular DNA structures were manually counted in a blinded manner as NET-positive tiles.

### Live fluorescence microscopy of enzymatic digest of human NETs

500 μL of 1.6 × 10^6^ cells/mL of human blood PMNs were seeded in a 4-well glass bottom microscopy cell culture dish (Greiner, 627871) and rested for 30 min before stimulation at 37°C and 5% CO2 with PMA (600 nM) for 3 h. After stimulation, the medium was carefully removed and the cells were washed with PBS before adding fresh culture medium (RPMI culture medium without phenol red (Sigma-Aldrich, R7509) ⍰.+ ⍰10% FBS (heat inactivated, sterile filtered, TH Geyer, 11682258)). Hoechst 33342 (Thermo Fisher, 1 μg/mL) to stain nuclear DNA and SYTO RNAselect Green fluorescent dye (Thermo Fisher, 50 µM) to stain naRNA was added to the cells, as well as RNase A (Thermo Fisher, EN0531; 100 μg/mL) was added between time point 0 and 5 min. Live cell imaging was performed using a Zeiss LSM800 Confocal microscope with a 63x objective and z-stack acquisition, taking an image every 5 min for 30-60 min respectively. Image analysis and video creation was performed using ImageJ-Win64.

### Click chemistry of primary, stem cell-derived PMNs and fluorescence microscopy

500 μL of 1.6 × 10^6^ cells/mL of human stem cell-derived PMNs treated with 5-ethynyluridine or left untreated were seeded in a 24-well plate containing poly-L-lysine-coated glass coverslips and rested for 30 min before stimulation with PMA (600 nM) at 37°C and 5% CO_2_ for 12 h. After stimulation, the cells were washed and permeabilized with ice-cold acetone (Applichem, A1582.2500PE) 1:1 methanol (Honeywell, 32213-2.5L) for 5 min at RT. Subsequently, the click chemistry (reagents see Supplementary Table S4) labeling of endogenous RNA was performed as described (*18*). Briefly, in a total volume of 500 μL, 2 μL of AF546-Azide, a pre-mixture of 1 mM CuSO_4_ and 1.25 mM THPTA, 5 mM aminoguanidine-hydrochloride and 5 mM Na-ascorbate in PBS were added to the cells in a 24-well plate. The wells were sealed with plastic foil and incubated for 1 h at RT while shaking in the dark. For negative controls, 5-ethynyluridine untreated cells incubated with complete click chemistry reagents and 5-ethynyluridine treated cells incubated with PBS and AF546-Azide only were used. No significant signals were observed. After the incubation, the cells were washed three times for 5 min with PBS and counterstained with rRNA Y10b-AF647 (see Supplementary Table S3) at 1:50 in PBS for 2 h at RT in the dark. After repeated washing, the cells were incubated with Hoechst 33342 to stain nuclear DNA and mounted as described above. Imaging and analysis were performed as previously described.

### Electron microscopy

For electron microscopy, 500 μL of 1.6 × 10^6^ cells/mL of human blood-derived PMNs were seeded in a 24-well plate containing coverslips which were pre-coated with 0.01 % poly-L-lysine (Sigma, A-005-C) for 15 min at 37°C for scanning electron microscopy (SEM). For transmission electron microscopy (TEM), coverslips were coated with a carbon layer, glow-discharged and UV-treated overnight before cells were seeded. Cells were rested for 30 min at 37°C and 5% CO_2_ and subsequently stimulated with 1200 nM PMA or left untreated for 3 h. Afterwards, the cells were fixed in 2.5% glutaraldehyde in PBS for 2 h at room temperature followed by incubation at 4°C. For SEM, samples were post-fixed with 1% osmium tetroxide for 1 h on ice. Subsequently, samples were dehydrated in a graded ethanol series followed by critical point drying (CPD300, Leica Microsystems) with CO_2_. Finally, the cells were sputter-coated with a 4 nm thick layer of platinum (CCU-010, Safematic) and examined with a field emission scanning electron microscope (Regulus 8230, Hitachi High Technologies) at an accelerating voltage of 3 kV. For antibody labeling, cells treated as described above were fixed in 4% formaldehyde in PBS for 1-2 hours at room temperature and 4°C overnight. After washing and blocking (0.2% gelatin in PBS) samples were incubated for 1 h at room temperature with rRNA Y10b as the primary antibody in blocking buffer and for 1 h at RT with goat anti-mouse antibodies coupled to 6 nm gold in blocking buffer (Jackson ImmunoResearch, code number 115-195-166). Samples labeled with 6 nm gold were further silver enhanced. Following immunolabeling, the samples were treated with 1% uranyl acetate for 5 min at RT, dehydrated and critical point dried as before. Samples were sputter-coated with a 5 nm thick layer of carbon (CCU-010, Safematic) and analyzed in the SEM with an accelerating voltage of 5 kV. For TEM, samples were prepared by the PLT method as described above for IF of ultrathin sections. Samples were placed on grids (Cu, Ni) and subsequently sectioned into 50-60 nm ultrathin slices. Immunolabeling was performed by incubation with anti-rRNA Y10b as the first primary antibody in blocking buffer at 4°C overnight, followed by incubation with anti-LL37 as the second primary antibody for 1 h at RT. Afterwards, the cells were incubated with goat anti-mouse coupled to 6 nm gold and goat anti-rabbit 12 nm gold secondary antibodies in blocking buffer for 1 h at RT. Following immunolabeling, the samples were treated with 1% uranyl acetate for 5 min at RT before analysis in the TEM.

### Transient transfection of HEK293T cells

HEK293T cells were transiently transfected with the respective TLR8, TLR7, TLR9 and NF-κB reporter plasmids as described (*20*) (see Supplementary Table S5 for plasmids) using X-tremeGENE™ HP DNA Transfection Reagent (Merck, 6366236001). A total amount of 5 × 10^4^ cells/well were seeded in a 24-well plate one day prior to transfection. For the transfection of one well, 100 ng of the according TLR plasmid, 100 ng of the firefly luciferase NF-κB reporter and 10 ng *Renilla* luciferase control reporter was mixed in Opti-MEM™ Reduced Serum Medium (Thermo Fisher, 31985062) in a total volume of 50 μL. After 15 min incubation at RT, the transfection mix was added to the cells and the cells were incubated for 48 h. Prior to subsequent stimulation, the medium was changed to complete DMEM medium (Sigma, D5796-24X500ML), and the cells were incubated with the respective NET content stimuli and controls (R848 (2.5 μg/mL), TL8 (100 ng/mL), CpG (1.25 μM) + DOTAP (25 μg/mL), ssRNA (0.6 μg/mL) + DOTAP (20 μg/mL), mock NETs + RNase inhibitor (1:50 dilution), PMA NETs (1:50 dilution), PMA NETs + RNase inhibitor (1:50)) for 18 h at 37°C and 5% CO_2_. Supernatants were then removed, and the cells frozen briefly at -80 °C. Subsequently they were used for dual luciferase assay.

### Dual luciferase reporter assay

The dual luciferase reporter assay for detection NF-κB activation after TLR transfection and subsequent stimulation was performed as described (*8*). In brief, supernatants were removed from the cells after stimulation and 60⍰.µL/well of 1x passive lysis buffer (E194A, Promega) was added. The plate was then incubated for 15⍰.min at RT on the plate shaker and subsequently stored at −80°C for at least 15⍰min to facilitate complete cell lysis. After thawing, the cell solution (60 μL) was transferred into a V-bottom 96-well plate and centrifuged for 10⍰min at 2500⍰.rpm and 4°C to pellet cell debris. Ten microliters of supernatant were then transferred into a white microplate and each condition was measured in triplicates using the FLUOstar OPTIMA device (BMG Labtech). Firefly and Renilla luciferase activity were determined using the Promega Dual luciferase kit. Both enzyme activities were measured for 12.5⍰s with 24 intervals of 0.5⍰s, respectively. The data was analyzed by calculating the ratio of the two measured signals, thereby normalizing each firefly luciferase signal to its corresponding Renilla luciferase signal. The ratios were represented as the relative light units (RLU) of NF-κB activation.

### Extracellular bacterial killing of *S. aureus*

The killing assay of *S. aureus* with human PMNs was performed according to Brinkmann *et al*., 2004 (*1*). In brief, PMNs were seeded at a density of 2 × 10^6^ cells/mL and incubated with PMA (600 nM) for 2 h at 37°C. Afterwards, the medium was carefully replaced with serum-free culture medium, containing 2% heat-inactivated pooled human serum with cytochalasin D (10 µg/mL) and incubated further for 15 min before infection with bacteria. Cytochalasin D treatment did not affect NETs and this concentration was effective in blocking bacterial phagocytosis. To investigate whether naRNA of NETs was important in extracellular killing, samples were either treated with RNase A (Thermo Fisher, EN0531; 100 μg/mL) or DNase I (Thermo Fisher, EN0521; 1 U/10 μL) for 2 h during NET formation, or after NET formation during the 30 min bacterial killing process. Samples were centrifuged at 700 x g for 10 min and incubated at 37°C and 5% CO_2_ for 30 minutes. Bacterial killing was measured as percentages of control values (bacteria incubated alone in media without neutrophils).

### In vivo analysis of naRNA DAMP effects

To investigate the effect of NETs with or without RNase inhibitors and the respective TLR signaling, 20 μL of NET content and the respective controls were injected into the ears of C57BL/6 and *Tlr13*^-/-^ mice intradermally on day 0. Afterwards, as a measure of inflammation, the ear thickness was assessed using a manual caliper (0.01–10 mm, Peacock, Tokyo, Japan) until day 4.

### Neutrophil infiltration *in vivo* fluorescence imaging

The *in vivo* experiment for investigating neutrophil infiltration was performed as described (*8*). Briefly, LysM^EGFP/+^ mice were injected intradermally with 20⍰µL of PMA or mock NET content or respective controls. LysM^EGFP/+^ mice were then anesthetized with inhalation isoflurane and *in vivo* fluorescence imaging was performed using the IVIS Lumina II imaging system (Caliper). EGFP fluorescence was measured using excitation (465⍰nm), emission (515–575⍰nm), and exposure time (0.5⍰s). Data are quantified as total radiant efficiency ([photons/s]/[µW/cm^2^]) within a circular region of interest using Living Image software (Caliper).

### Imiquimod model of psoriatic skin inflammation

To analyze the effect of RNA signaling in an *in vivo* model for psoriasis, the well-established imiquimod mouse model was used (*31*). C57BL/6 and *Tlr13*^-/-^ mice were used and, briefly, 70 μL (62.5 mg) of imiquimod (5%, Taro Pharmaceuticals Industries, Hawthorne, NY) was applied daily to both sides of a mouse ear for 5 consecutive days (day 0 to 4). Ear thickness was measured with a manual caliper (0.01–10 mm, Peacock, Tokyo, Japan) before imiquimod application. A day after the last application of imiquimod (day 5), full-thickness ear skin was excised with surgical scissors for histologic analysis.

### Histologic analysis of mouse ear skin

Ear skin specimens were collected by excising the entire ear with surgical scissors, fixed in formalin (10%), and embedded in paraffin. 4 μm thickness sections were mounted onto glass slides and stained with H&E by the Johns Hopkins Reference Histology Laboratory. To measure epidermal thickness, 100 epidermal thickness measurements per mouse were averaged from images taken at ×200 magnification (Hamamatsu Nanozoomer) using ImageJ software (NIH).

### ELISA

To measure cytokine release of BlaER-1, THP-1, N/TERT-1, NHEK, NK-92 MI, PBMCs and 3D human skin equivalent after stimulation with NET content and respective controls, ELISA Kits for hIL-8 (ELISA MAX™ Deluxe Set Human IL-8, Biolegend, 431504), hIL-6 (ELISA MAX™ Deluxe Set Human IL-6, Biolegend, 430504), IFN-γ (ELISA MAX™ Deluxe Set Human IFN-γ, Biolegend, 430104), and TNF (TNF alpha Human Uncoated ELISA Kit, Invitrogen, 88-7346-88) were used according to the manufacturer’s instructions. Samples were assessed in triplicates.

### qPCR analysis of IL-8 expression of 3D human skin equivalent

To investigate IL-8 expression of 3D human skin equivalent after stimulation with NET contents, qPCR analysis was performed. First, total RNA was isolated using the RNeasy Mini Kit (Qiagen, 74106) for animal tissues and cells. For cDNA preparation, the High-Capacity RNA-to-cDNA-Kit (Thermo Fisher, 4387406) was used according to the manufacturer’s instructions. For the qPCR, the TaqMan™ system was used. Briefly, a master mix of TaqMan™ Universal Mastermix II (Thermo Fisher, 4440040) and TaqMan™ Gene Expression Assay (Thermo Fisher, 4448892) was prepared according to the manufacturer’s instructions. For one reaction, 5.5 μL master mix and 4.5 μL of the respective cDNA (IL-8 and TBP) were mixed and the qPCR was run. Analysis in triplicates was performed using QuantStudio Real-Time-PCR software version 1.3 (Thermo Fisher).

### Statistics

Experimental data were analyzed using Excel 2019 (Microsoft) and/or GraphPad Prism 8, microscopy data with ImageJ-Win64 or ZenBlue3 software, flow cytometry data with FlowJo V10. Normal distribution in each group was always tested using the Shapiro–Wilk test first for the subsequent choice of a parametric (ANOVA, Student’s t-test for normally distributed data) or non-parametric (Mann–Whitney U) test as indicated in figure legends. p-values (α = 0.05) were then calculated, and multiple testing was corrected for in Prism, as indicated in the figure legends. Values < 0.05 were generally considered statistically as significant and denoted by * throughout even if considerably lower. Comparisons made to unstimulated control, unless indicated otherwise, were denoted by brackets.

## Supplementary figures

**Fig. S1: Controls of IF microscopy, stem cell-derived PMNs, and electron microscopy**

**(A)** Confocal microscopy of unstimulated primary human PMNs (control to Fig. 1A) after 3 h and stained for naRNA (anti-rRNA Y10b, magenta) and DNA (Hoechst 33342, white, n = 3, representative images, scale bar: 10 μm). **(B)** Confocal microscopy of unstimulated primary murine BM-PMNs (control to Fig. 1C) after 16 h and stained as in A (n = 3, representative images, scale bar: 10 μm). **(C)** Brightfield microscopy analysis of cytospun control primary human stem cell-derived PMNs (control to Fig. 1D) differentiated *in vitro* with/without 100 μM 5-Ethynyluridine (5-EU, n = 3, representative images, scale bar: 20 μm). **(D)** FACS analysis for cells shown in **C** and Fig. 1D (n=3, representative data). **(E)** Scanning electron microscopy of PMA-treated human primary PMNs showing only secondary antibody staining (no primary antibody) control of Fig. 1F (n = 1, representative data; the image on the right is a composite image with signals from secondary electron and backscattered electron detectors for topography and additional material information, respectively).

**Fig. S2: Antibacterial effect of NETs on live S. aureus, controls of IF microscopy and line plot analysis of naRNA and LL37**

**(A)** Extracellular bactericidal activity of human PMNs/NETs after infection with *S. aureus* and treatment with RNase A and DNase I during or after formation of PMA-induced NETs (n = 8, combined data, mean+SD, *p<0.05 according to one-way ANOVA). **(B)** Confocal microscopy of unstimulated or PMA-stimulated primary human PMNs (control to Fig. 2B) after 3 h, stained DNA (Hoechst 33342, white, n = 9, representative images, scale bar: 10 μm). **(C)** as in **B** but controls of Fig. 2D (n = 3). **(D)** As in **B** but controls of Fig. 2F (n = 3).

**Fig. S3: Inhibition of PAD4 in human PMNs during NET formation assay and *Unc93b1***^***-/-***^ **and *Tlr9***^***-/-***^ **BM-PMN stimulation with human NETs**

**(A)** Confocal microscopy of stimulated primary human PMNs after 3 h and stained for DNA (Hoechst 33342, white) in the presence or absence of the PAD4-inhibitor Cl-amidine (200 μM, n = 3, representative images; scale bar 10 μm). **(B)** Confocal microscopy of primary C57BL/6 WT or *Unc93b1*^-/-^ murine BM-PMNs stimulated as indicated for 16 h as in A (n = 3 WT, n = 1 *Unc93b1*^-/-^, representative images, scale bar: 10 μm). **(C)** Confocal microscopy of primary C57BL/6 WT or *Tlr9*^-/-^ murine BM-PMNs stimulated as indicated for 16 h as in **A** (n = 3, representative images, scale bar: 10 μm).

**Fig. S4: Immune responses of PBMCs, macrophages, and NK-cells to NETs and in vivo fluorescence imaging**

**(A)** Triplicate IL-8 ELISA from WT, *TLR8*^*-/-*^ and *TLR7*^*-/-*^ THP-1 cells stimulated as indicated for 18 h, normalized to PMA+ionomycin control (n = 4, combined data, mean+SD, each dot represents one biological replicate, *p<0.05 according to one-way ANOVA). **(B-D)** Triplicate ELISA for TNF (**A**, n = 4), IL-6 (**B**, n = 3), and IL-8 (**C**, n = 3) from primary human PBMCs stimulated as indicated with/without CU-CPT9a for 24 h (combined data, mean+SD, each dot represents one biological replicate, *p<0.05 according to one-way ANOVA). **(E)** Triplicate IFN-γ ELISA from NK-92 MI cells stimulated as indicated for 24 h (n = 3, combined data, mean+SD, each dot represents one biological replicate, *p<0.05 according to one-way ANOVA).

**Fig. S5: Pre-association of naRNA and LL37 in resting healthy donor neutrophils**

**(A)** Confocal microscopy of primary human PMNs stimulated as indicated for 3 h and stained for naRNA (anti-rRNA Y10b, magenta), LL37 (anti-hLL37-DyLight550, yellow) and DNA (Hoechst 33342, white, n = 3, representative images). The line plot analysis of LL37, RNA and DNA staining was performed using ZenBlue3 software. Two to three different line plots from the same representative image are shown. **(B)** Transmission electron microscopy of unstimulated human primary PMNs using anti-rRNA and anti-hLL-37 primary and immunogold (6 nm (black arrow) and 12 nm (white arrow) respectively)-labeled secondary antibodies (n = 3, representative images, scale bars as indicated).

**Fig. S6. Graphical abstract**

## Supplemental movies

**Supplemental movie S1: 3D reconstruction of NET DNA network decorated with naRNA**

Confocal microscopy of primary human PMNs stimulated with PMA for 3 h, stained for naRNA (anti-rRNA Y10b, magenta) and DNA (Hoechst 33342, white, n = 3) and 3D analysis and animation were performed using ZenBlue3 software.

**Supplemental movie S2: Digest of naRNA in PMA-induced NETs**

Confocal live microscopy of RNase digestion of NETs obtained from primary human PMNs stimulated with PMA for 3 h, stained for naRNA (SYTO RNAselect, magenta) and DNA (Hoechst 33342, white, n = 3). RNase A was added between time point 0 and 5 min (n = 3, scale bar 20 μm). Video analysis and animation were performed using ImageJ-Win64 software.

## Supplementary Tables

**Supplementary Table 1:**
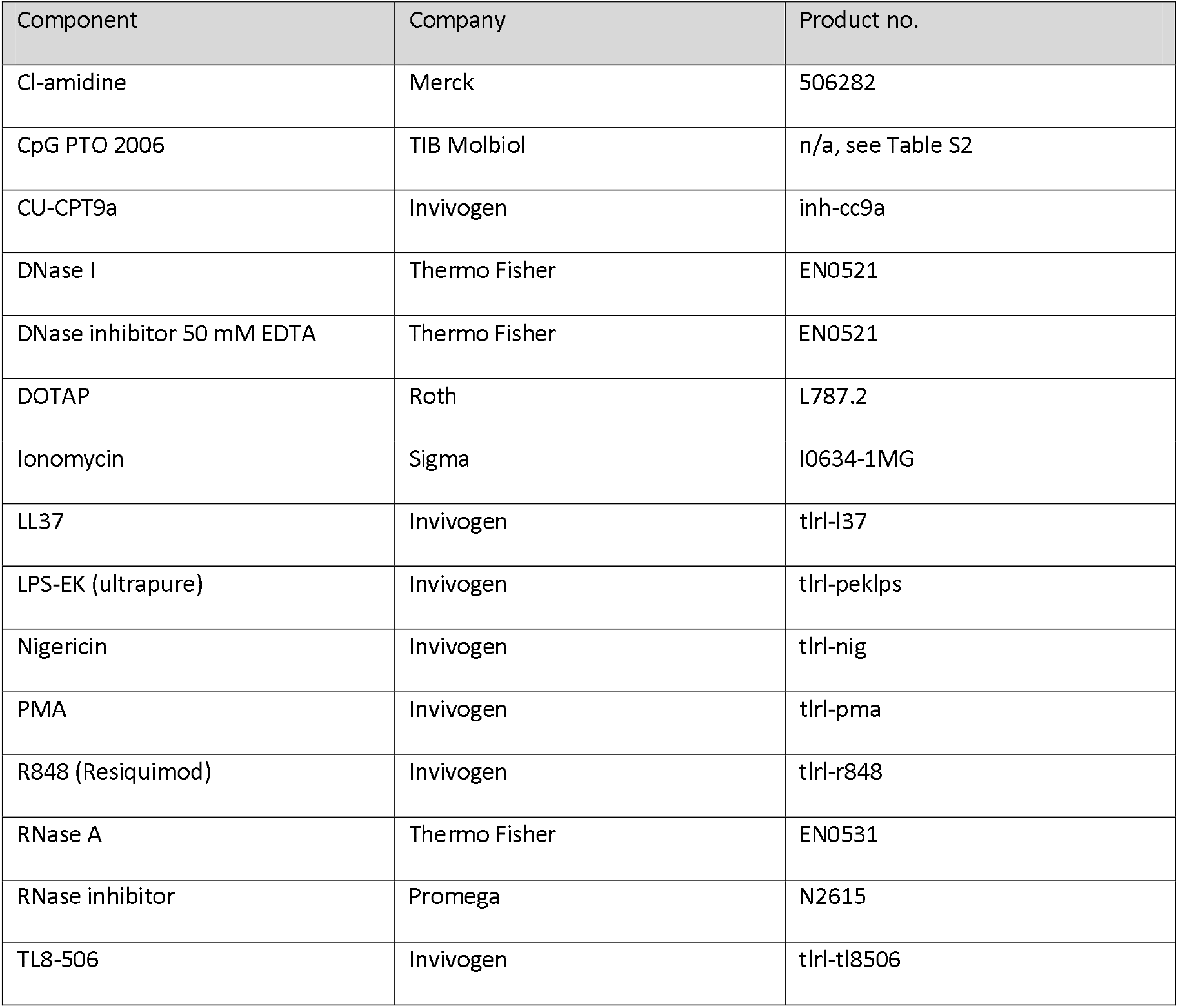
commercial TLR ligands and inhibitors; enzymes.

**Supplementary Table 2:**
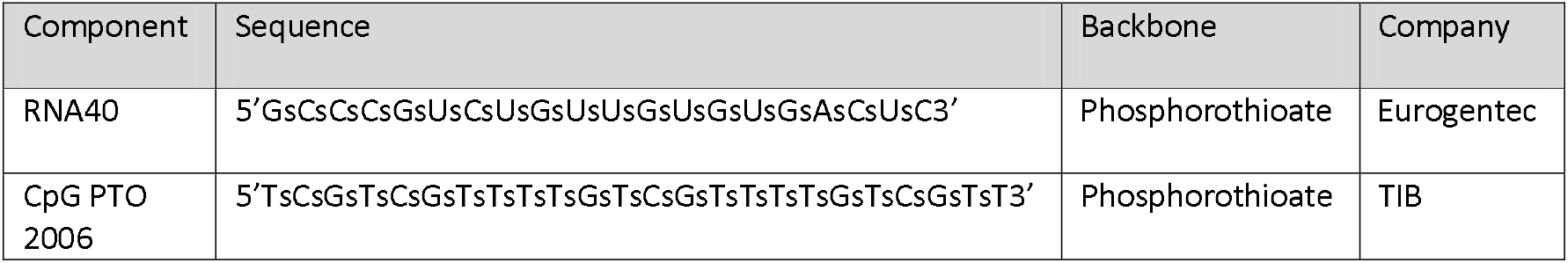
Nucleic acid TLR agonists.

**Supplementary Table 3:** Antibodies and conjugation kit.

| Item | Fluorophore | Species | Isotype | Company | Product no. |
| --- | --- | --- | --- | --- | --- |
| Anti-hLL37 | DyLight 550 | rabbit | IgG | LSBio | LS-B6696-500 |
| DyLight 550 Conjugation Kit (Fast) | DyLight 550 | - | - | Abcam | ab201800 |
| Anti-rRNA (Y10b) | unconjugated | mouse | IgG<sub>2a</sub> K | Santa Cruz Biotechnology | sc-33678 |
| Anti-rRNA (Y10b) Alexa Fluor® 647 | AF647 | mouse | IgG<sub>2a</sub> K | Santa Cruz Biotechnology | sc-33678<br>AF647 |
| Anti-mouse IgG | AF647 | chicken | IgY | Thermo Fisher | A-21463 |
| Hoechst 33342 | - | - | - | Thermo Fisher | H21492 |
| SYTO RNAselect Green fluorescent dye | - | - | - | Thermo Fisher | S32703 |
| Anti-mouse Cy3 | Cy3 | goat | IgG | Jackson ImmunoResearch | 115-165-146 |
| Anti-rabbit AF488 | AF488 | goat | IgG | Molecular Probes | A-11008 |
| Anti-mouse 6 nm gold | - | goat | IgG | Jackson ImmunoResearch | 115-195-166 |
| Anti-rabbit 12 nm gold | - | goat | IgG | Jackson ImmunoResearch | 111-205-144 |

**Supplementary Table 4:**
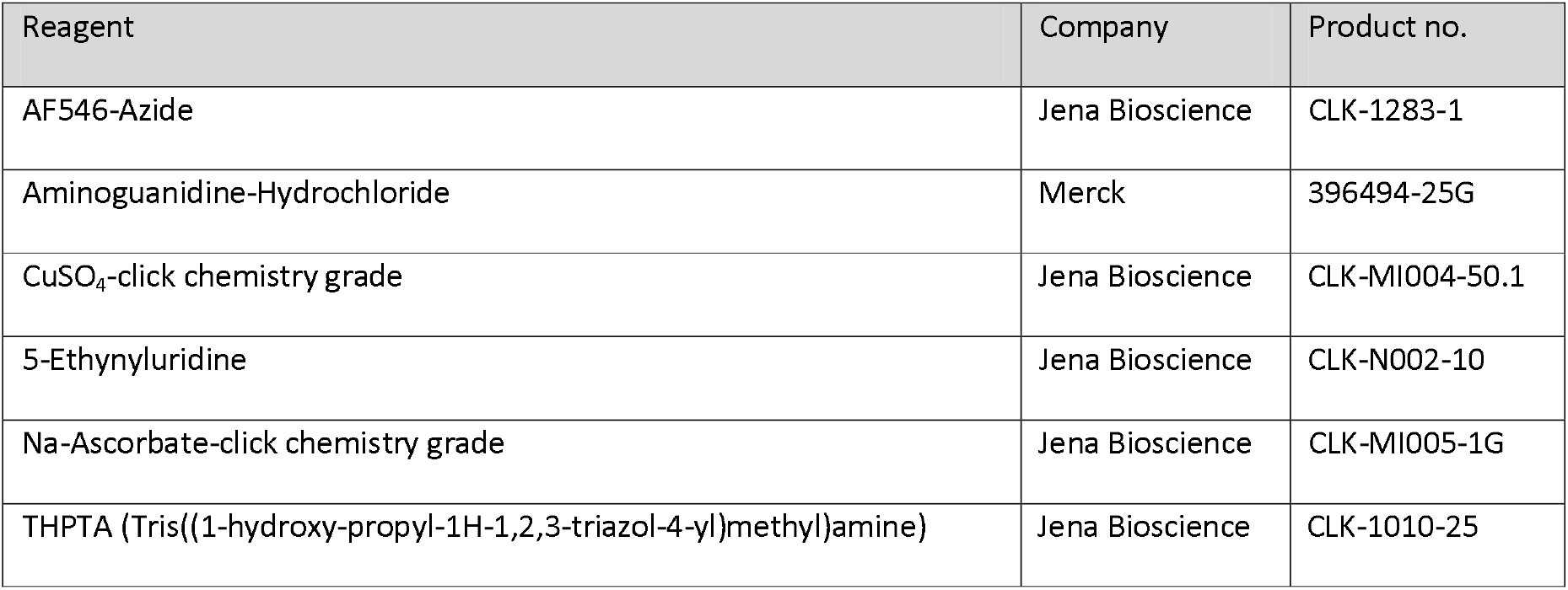
Click chemistry reagents.

| Reagent | Company | Product no. |
| --- | --- | --- |
| AF546-Azide | Jena Bioscience | CLK-1283-1 |
| Aminoguanidine-Hydrochloride | Merck | 396494-25G |
| CuSO<sub>4</sub>-click chemistry grade | Jena Bioscience | CLK-MI004-50.1 |
| 5-Ethynyluridine | Jena Bioscience | CLK-N002-10 |
| Na-Ascorbate-click chemistry grade | Jena Bioscience | CLK-MI005-1G |
| THPTA (Tris((1-hydroxy-propyl-1H-1,2,3-triazol-4-yl)methyl)amine) | Jena Bioscience | CLK-1010-25 |

**Supplementary Table 5:** Plasmids used for HEK293T transfection.

| Plasmid name insert | Vector backbone | Insert |
| --- | --- | --- |
| NF-κB reporter | pGL3 | 6x NF-κB response element |
| Renilla | pRL-TK | *Renilla* |
| hTLR7 | pcDNA3.1 (+) | hTLR7 |
| hTLR8 | pcDNA3.1 (+) | hTLR8 |
| hTLR9 | pSEM-hTLR9 | hTLR9 |

**Supplementary Table 6:** primers used for qPCR.

| Gene | Assay number |
| --- | --- |
| IL-8 | Hs00174103_m1 |
| TBP (housekeeper) | HS00427620_m1 |

**Supplementary Table 7:** 10x Ammonium chloride erythrocyte lysis buffer.

| Compound | Company | Product no. |
| --- | --- | --- |
| 1.54 M NH<sub>4</sub>Cl | Roth | 5470.1 |
| 100 mM KHCO<sub>3</sub> | Fluka | 60220 |
| 1 mM EDTA; pH 8 | Thermo Fisher | 15575020 |
| dissolved in Ampuwa water | Fresenius Kabi | 1833 |
| pH adjusted to 7.3, sterile filtered (0.22 µm) |  |  |

