## Supplementary figures and images for "Release of the pre-assembled naRNA-LL37 composite DAMP re-defines neutrophil extracellular traps (NETs) as intentional DAMP webs"

### Supplementary Figure S1

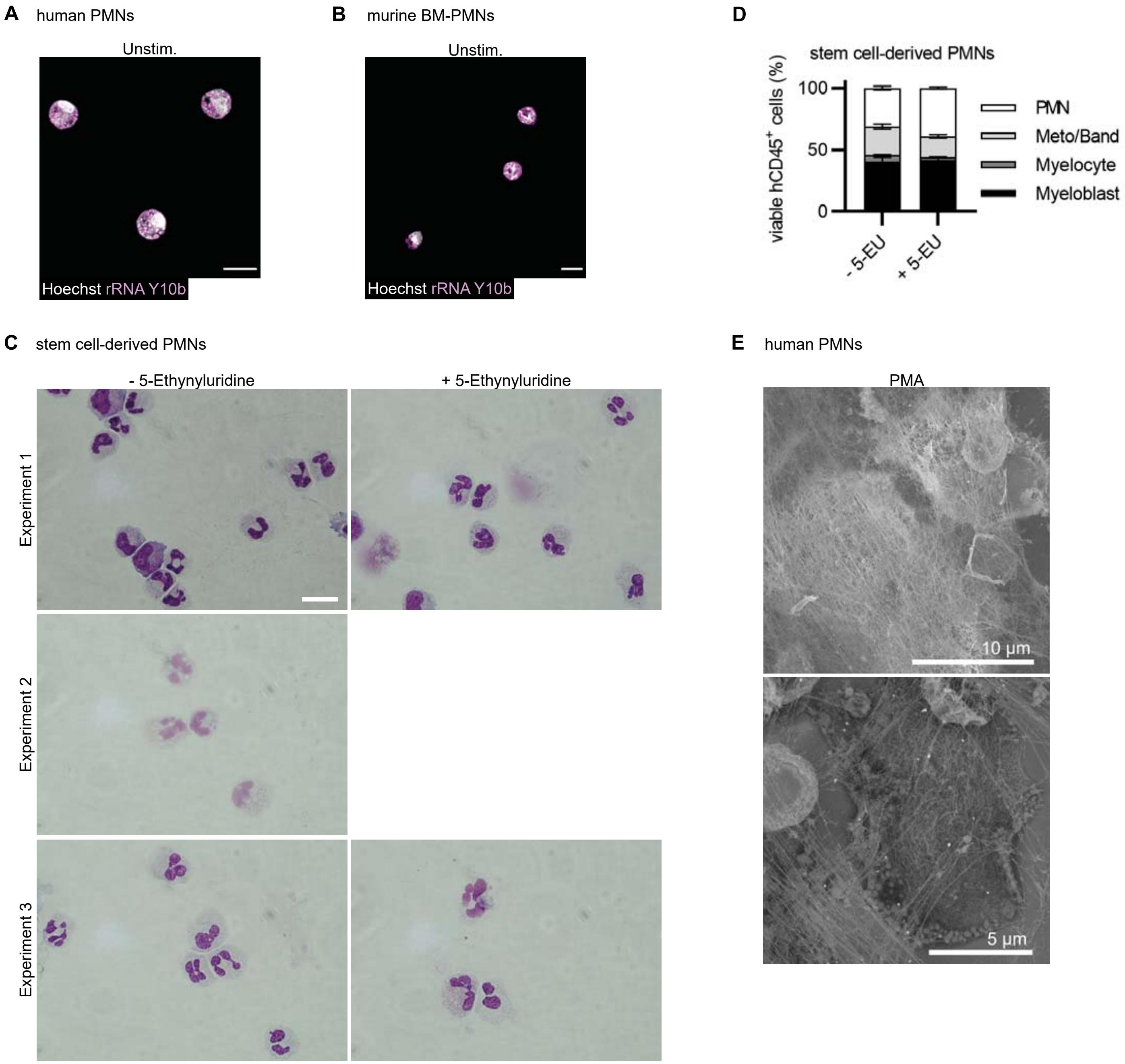

### Supplementary Figure S2

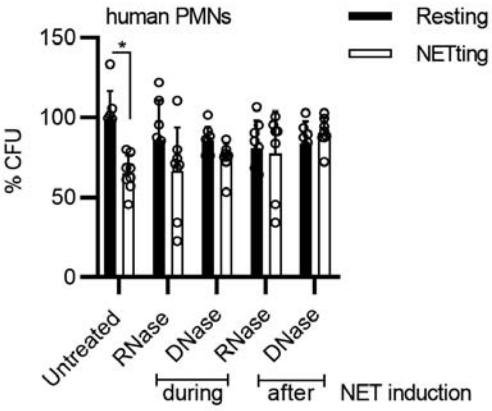

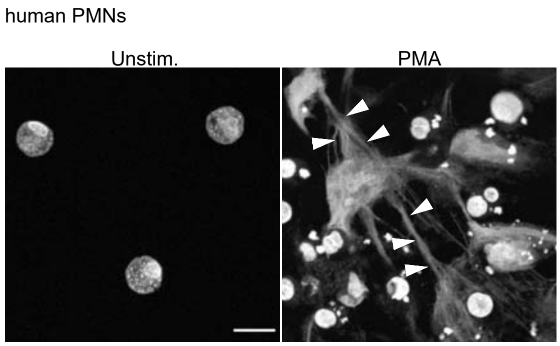

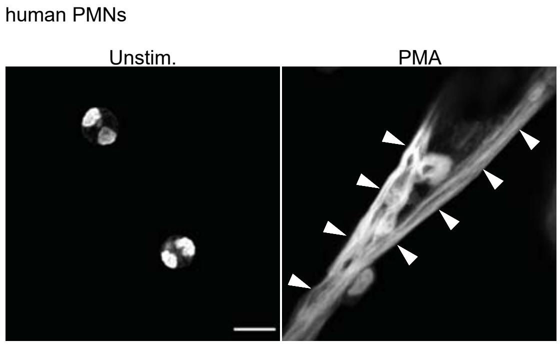

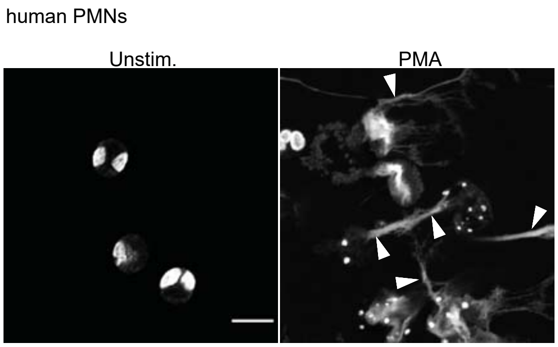

### Supplementary Figure S3

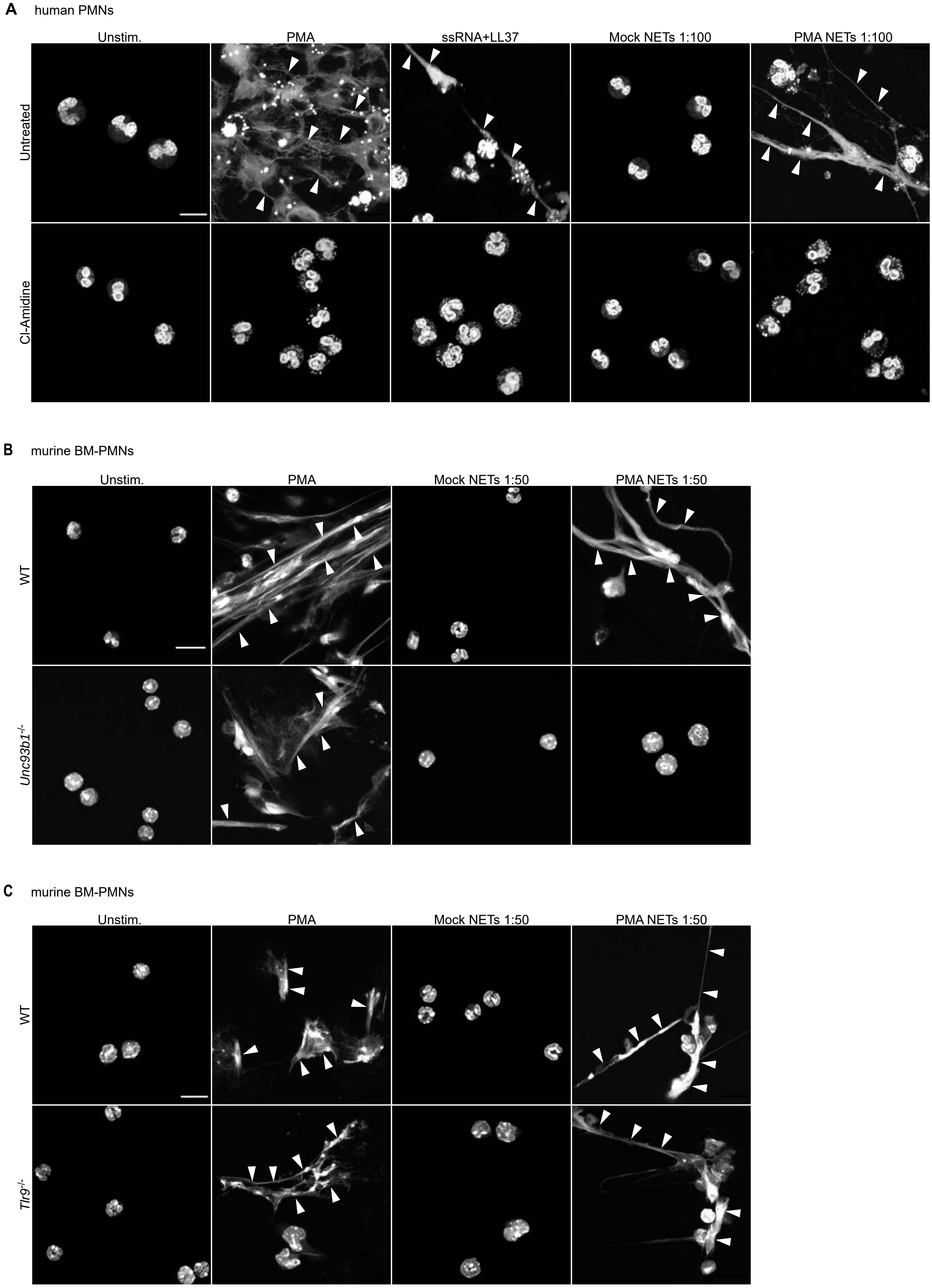

### Supplementary Figure S4

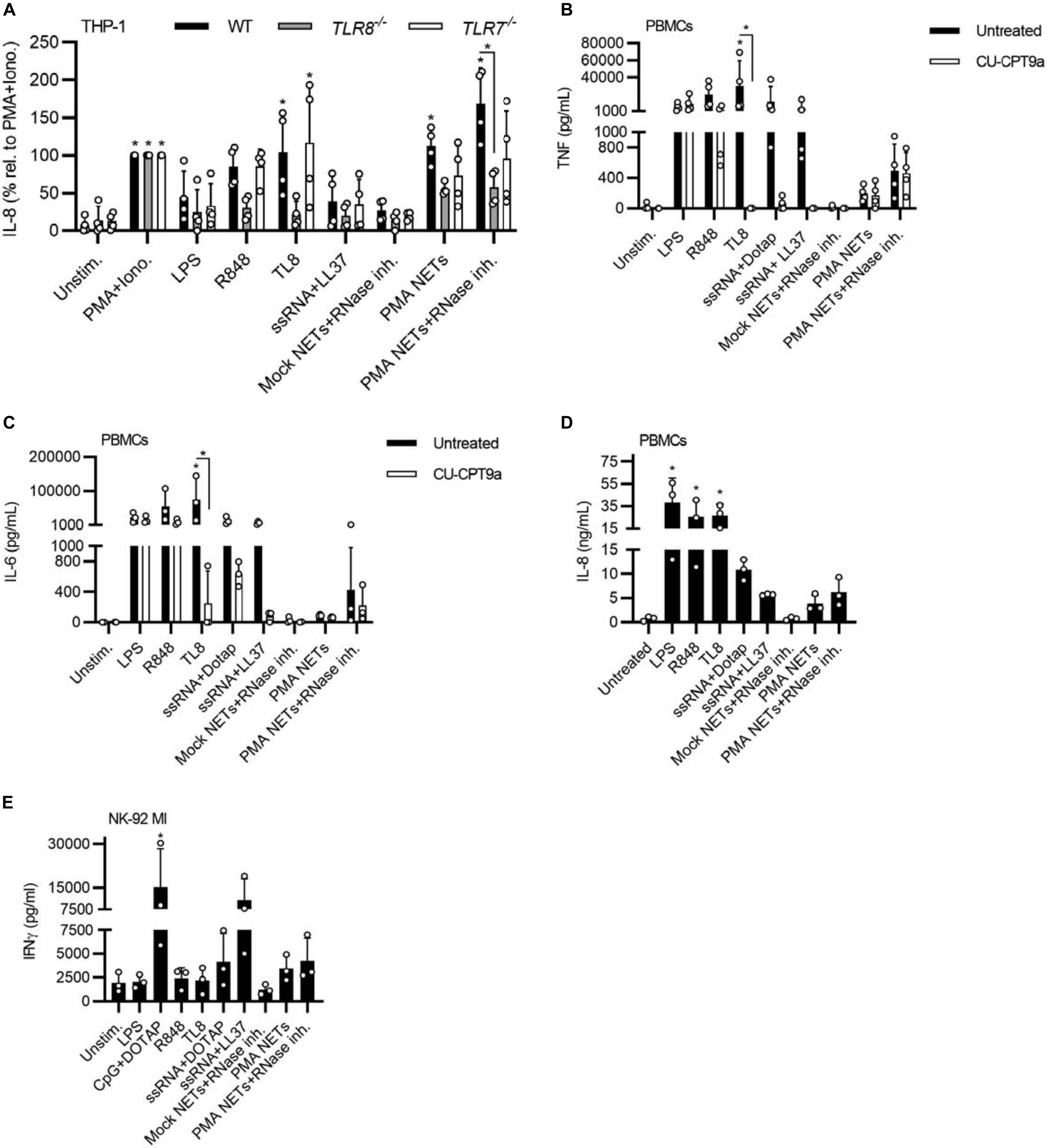

### Supplementary Figure S5

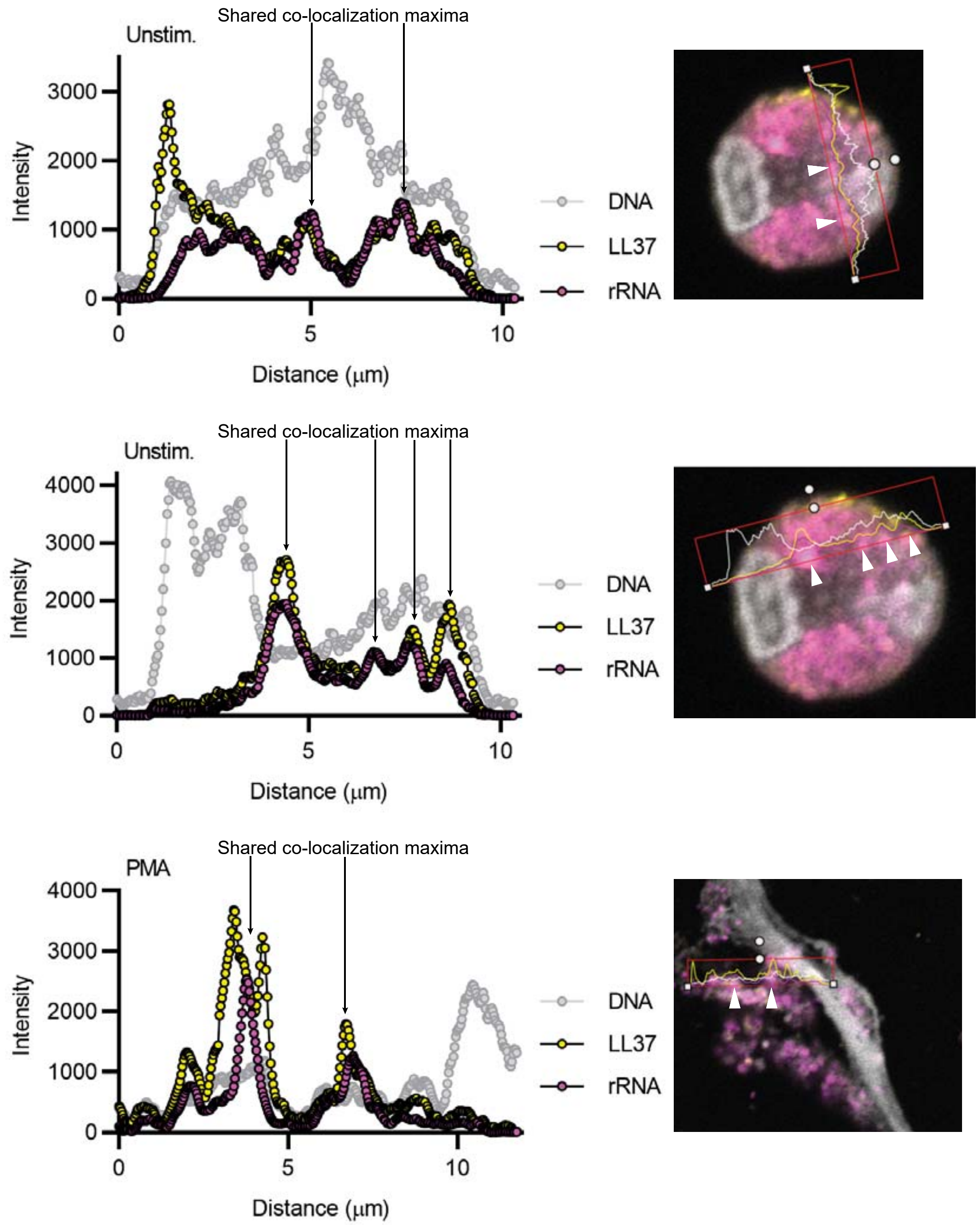

### Supplementary Figure S6

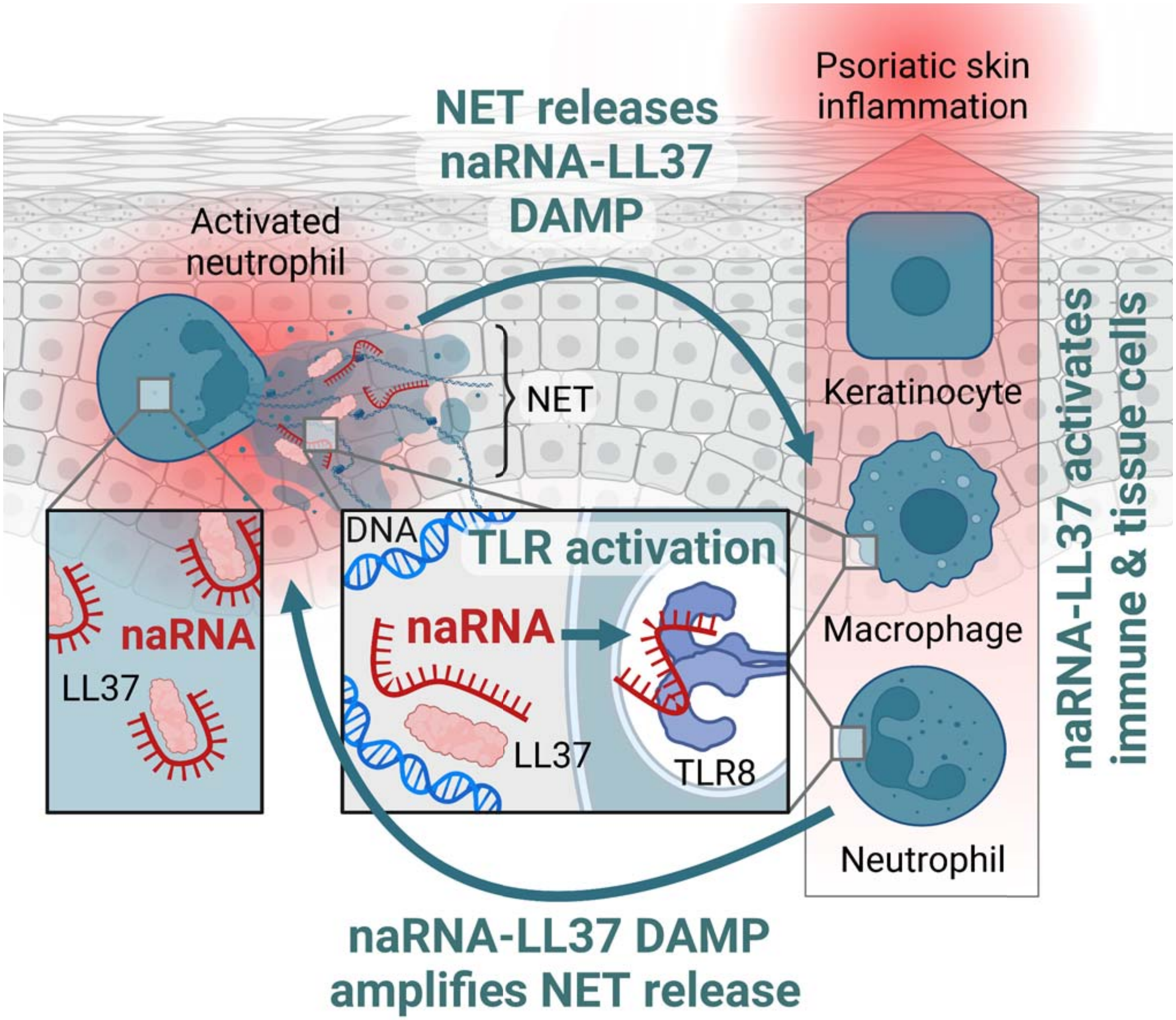
